# *In vitro* evolution of *Listeria monocytogenes* reveals selective pressure for loss of SigB and AgrA function at different incubation temperatures

**DOI:** 10.1101/2022.02.23.481730

**Authors:** Duarte N. Guerreiro, Jialun Wu, Emma McDermott, Dominique Garmyn, Peter Dockery, Aoife Boyd, Pascal Piveteau, Conor P. O’Byrne

## Abstract

The alternative sigma factor B (σ^B^) contributes to the stress tolerance of the foodborne pathogen *Listeria monocytogenes* by upregulating the General Stress Response. We previously showed that σ^B^ loss-of-function mutations arise frequently in strains of *L. monocytogenes*, and suggested that mild stresses might favour the selection of such mutations. In this study, we performed *in vitro* evolution experiments (IVEE) where *L. monocytogenes* was allowed to evolve over 30 days at elevated (42°C) or lower (30°C) incubation temperatures. Isolates purified throughout the IVEE revealed the emergence of *sigB* operon mutations at 42°C. However, at 30°C independent alleles in the *agr* locus arose, resulting in the inactivation of the Agr quorum sensing. Colonies of both *sigB*^−^ and *agr*^−^ strains exhibited a greyer colouration on 7-days-old agar plates compared with the parental strain. Scanning electron microscopy revealed a more complex colony architecture in the wild type than in the mutant strains. *sigB*^−^ strains outcompeted the parental strain at 42°C, but not at 30°C, whilst *agr*^−^ strains showed a small increase in competitive fitness at 30°C. Analysis of 40,080 *L. monocytogenes* publicly available genome sequences revealed a high occurrence rate of premature stop codons in both the *sigB* and *agrCA* loci. An analysis of a local *L. monocytogenes* strain collection revealed 5 out of 168 strains carrying *agrCA* alleles. Our results suggest that the loss of σ^B^ or Agr confer an increased competitive fitness in some specific conditions and this likely contributes to the emergence of these alleles in strains of *L. monocytogenes*.

**Importance:** To withstand environmental aggressions *L. monocytogenes* upregulates a large regulon through the action of the alternative sigma factor B (σ^B^). However, σ^B^ becomes detrimental for *L. monocytogenes* growth under mild stresses, which confer a competitive advantage to σ^B^ loss-of-function alleles. Temperatures of 42°C, a mild stress, are often employed in mutagenesis protocols of *L. monocytogenes* and promote the emergence of σ^B^ loss-of-function alleles in the *sigB* operon. In contrast, lower temperatures of 30°C promote the emergence of Agr loss-of-function alleles, a cell-cell communication mechanism in *L. monocytogenes*. Our findings demonstrate that loss-of-function alleles emerge spontaneously in laboratory-grown strains. These alleles rise in the population as a consequence of the trade-off between growth and survival imposed by the activation of σ^B^ in *L. monocytogenes*. Additionally, our results demonstrate the importance of identifying unwanted hitchhiker mutations in newly constructed mutant strains.

## Introduction

The Gram-positive bacterium *Listeria monocytogenes* is a robust foodborne pathogen found in a wide range of environments, such as soil, water, and faeces from both animals and humans (reviewed in (Hafner et al., 2021; NicAogáin & O’Byrne, 2016). *L. monocytogenes* is the aetiological agent of the disease listeriosis and is frequently associated with high mortality rates (typically 20-30%) (WHO, 2018). Listeriosis is especially hazardous for children, pregnant women, elderly people and immunocompromised patients. To establish an infection *L. monocytogenes* relies on its ability to withstand the extreme acidic pH of the stomach, as well as the osmotic stress and bile encountered in the duodenum (Gaballa et al., 2019; Sleator et al., 2009; Tiensuu et al., 2019).

The alternative sigma factor σ^B^ is responsible for the transcriptional control of a large regulon (~300 genes, or approximately 10% of *L. monocytogenes* genes) designated the general stress response (GSR) regulon, whose principal function is to protect against environmental stress (reviewed in (Guerreiro, Arcari, et al., 2020; Liu et al., 2019; Tiensuu et al., 2019). σ^B^ is also implicated in *L. monocytogenes* virulence by regulating the expression of the internalins *inlA* and *inlB* (Kim et al., 2005). These cell wall-associated proteins enhance *L. monocytogenes* invasion towards non-phagocytic cells at the intestinal epithelium (Ireton & Cossart, 1997; Lecuit et al., 2001) through interactions with the E-cadherin and Met receptors, respectively (Mengaud et al., 1996; Shen et al., 2000). The activity of σ^B^ is controlled through a complex signal transduction cascade, where a stress-sensing protein complex known as the stressosome (composed of RsbR, RsbS and RsbT) integrates environmental stress signals into the σ^B^ pathway. The current model proposes that following a stressful stimulus the kinase RsbT phosphorylates RsbR and RsbS leading to the subsequent release of RsbT from the stressosome. Free RsbT then interacts with and activates the phosphatase activity of RsbU, whose substrate is the phosphorylated anti-anti sigma factor RsbV. RsbV then sequesters RsbW, the anti-sigma factor that prevents σ^B^ from associating with RNA polymerase under non-stress conditions. The RsbV-RsbW interaction favours the association of σ^B^ with the RNA polymerase, to form the holoenzyme *E*σ^B^ and consequently transcription of the GSR regulon. Once the stress conditions abate, or the cell has adequately responded to them, the stressosome is reset to a sensing-competent state through the action of the RsbX phosphatase, which re-establishes the appropriate phosphorylation state of RsbR and RsbS (Oliveira et al., 2021; Xia et al., 2016). All the proteins involved in this signal cascade are encoded by the *sigB* operon, which comprises *rsbRSTUVW*-*sigB*-*rsbX* (Ferreira et al., 2004). σ^B^ plays an important role in enhancing *L. monocytogenes* resistance towards a wide range of lethal stresses (reviewed in (Guerreiro, Arcari, et al., 2020; NicAogáin & O’Byrne, 2016; O’Byrne & Karatzas, 2008). However, deploying σ^B^ under certain mildly stressful conditions comes with a cost, evidenced by an increase in growth rate for mutants lacking this sigma factor (Brøndsted et al., 2003; Chaturongakul & Boor, 2004; O’Donoghue et al., 2016). Recently we reported that mild heat stress (42°C) gives mutants lacking σ^B^ a competitive advantage in mixed cultures (Guerreiro, Wu, et al., 2020). Many of the genetic techniques used with *L. monocytogenes* rely on plasmids with a temperature-sensitive replication origin unable to replicate at temperatures above 37°C (e.g. *ori* pE194^ts^ in pMC39 a transposon delivery vector (Cao et al., 2007) and pMAD, a vector widely used for *L. monocytogenes* mutagenesis (Arnaud et al., 2004)). We also reported that the rate of premature stop codon (PMSC) occurrence in the *sigB* operon is unusually high, suggesting that sequenced strains might be subject to selective pressures for loss of σ^B^ function (Guerreiro, Wu, et al., 2020). Taken together these observations suggested that mild stress during laboratory culture might select for σ^B^ loss-of-function mutations. The present study sought to investigate whether extended growth at elevated temperature (42°C) would give rise to mutations affecting σ^B^ activity.

In *L. monocytogenes* quorum sensing is mediated by the Agr system, which is encoded by a four-gene operon (*agrBDCA*). AgrC (histidine kinase) and AgrA (response regulator) form a two-component system that senses the concentration of a signalling peptide that is encoded by *agrD* and secreted by AgrC (Zetzmann et al., 2016). The Agr system is crucial for cell surface attachment and biofilm formation when *L. monocytogenes* is growing at lower temperatures (25-30°C) (Garmyn et al., 2011; Riedel et al., 2009; Rieu et al., 2008). Deletion of either the *agrA* or *agrD* genes in *L. monocytogenes* EGD-e results in significant changes in global gene expression in log-phase cultures. There is a significant overlap between the genes affected by the loss of Agr and the σ^B^ regulon, suggesting a regulatory connection between the general stress response and quorum sensing systems (Dorey et al., 2019; Garmyn et al., 2012; Marinho et al., 2020; Riedel et al., 2009).

In this study, *in vitro* evolution experiments (IVEE) were performed to determine whether mutations affecting σ^B^ activity were selected in populations passaged for a 30-day period at 42°C compared to control cultures grown at 30°C. A colony phenotype associated with loss-of σ^B^ function was used to detect the emergence of potential *sigB* operon mutations in evolved populations; mutants lacking *sigB* have a grey colony phenotype (Guerreiro, Wu, et al., 2020). Whole genome sequencing, performed on multiple isolates from both populations, revealed that *sigB* loss-of-function mutations arise at 42°C but not at 30°C. However, grey colony variants did emerge at 30°C but were found to harbour mutations in the *agr* operon, specifically in *agrA* and *agrC*, but not in the *sigB* operon. An analysis of the published genome sequences of over 40,000 *L. monocytogenes* strains revealed that *agrA* and *agrC* mutations occur at a high rate compared to the other two-component systems suggesting a selective pressure. Our results show that loss-of-function mutations in both the *sigB* and *agr* operons can be selected during routine laboratory culture. Both the σ^B^ and Agr systems can influence colony architecture, as indicated by SEM analysis of the grey colony variants that arose at both temperatures. The study further highlights the importance of routine whole genome sequencing when studying phenotypic traits of *L. monocytogenes* isolates.

## Results

### IVEE performed at 42°C and 30°C promotes the emergence of grey coloured colonies

In our previous study, we demonstrated that *L. monocytogenes* strains with *sigB*^−^ genotypes and phenotypes exhibit increased growth rate and competitive advantage when grown in BHI at 42°C but not at 30°C (Guerreiro, Wu, et al., 2020). In this study, we sought to investigate the impact of incubation temperatures on the spontaneous emergence of alleles within the *sigB* operon. Cultures of *L. monocytogenes* EGD-e wild type strain were grown at 42°C in BHI for a total of 30 daily passages (see Material and Methods). The previously identified σ^B^-dependent grey colony colouration phenotype was used to distinguish potential *sigB*^−^ colonies from wild type (Fig. 1C) (Guerreiro, Wu, et al., 2020). Grey coloured colonies were detected from passage 15, at a relative abundance of 0.39% within the population and progressively increased to ~21% relative abundance in the following passages (Fig. 1A). Interestingly, grey coloured colonies were also detected in the IVEE performed at 30°C. These variants emerged from passage 10, from an initial relative abundance of ~0.1%, and increased to ~1% until the end of the experiment (Fig. 1B). The initial results suggested that *sigB*^−^ alleles emerge spontaneously and quickly increase in number within the parental population growing at both 42°C and 30°C temperatures, albeit to a greater extent at 42°C

**Figure 1 –.**
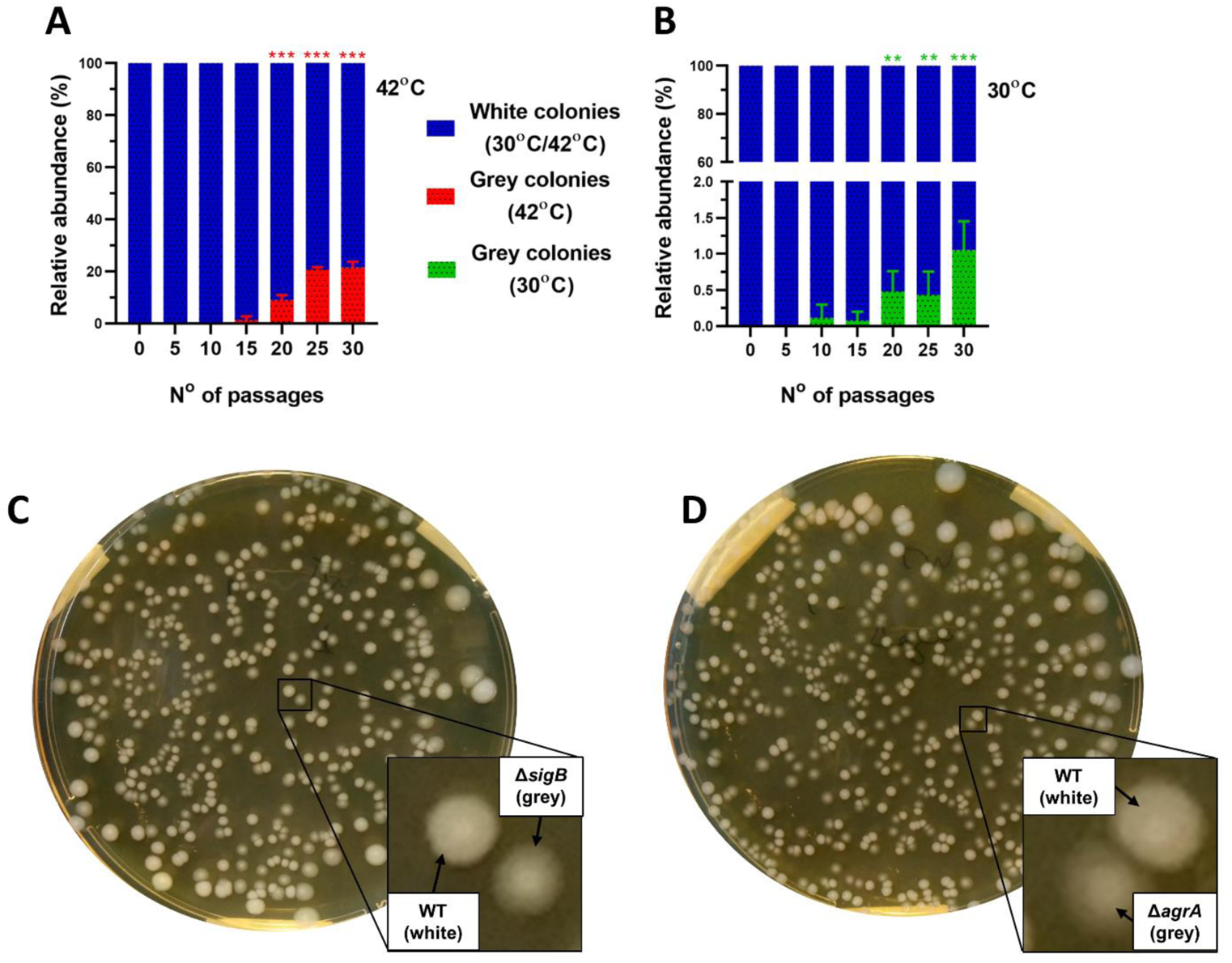
Differential emergence of grey-colony phenotypic variants during *in vitro* evolution experiments at 42°C and 30°C. *In vitro* evolution experiments where cultures of *L. monocytogenes* EGD-e wild type strain were grown in BHI at either (A) 42°C or (B) 30°C overnight. Passages were made every 24 hours for a total of 30 days by diluting the cultures in a 1:100 ratio into fresh BHI. Every 5 days the number of white and grey colonies was used to calculate the percentage of relative abundance. C and D show the colony colouration between the whiter WT and the greyish Δ*sigB* (C) and Δ*agrA* (D) strains. At least two independent biological replicates were performed for each incubation temperature. Statistical analysis was performed using a paired student *t*-test relative to passage 0 (**, *p*-value of <0.01; ***, *p*-value of <0.001).

### Mutations arise in the *sigB* operon at 42°C and in *agrCA* at 30°C

To identify genotype(s) responsible for the grey colony colouration observed from IVEE, we obtained whole genome sequence (WGS) from 14 colonies isolated from the IVEE performed at 42°C (9 grey and 5 white) and 10 colonies isolated from the IVEE performed at 30°C (6 grey and 4 white). We verified that all of the analysed grey coloured colonies isolated at 42°C carried a single nucleotide polymorphism (SNP) in an open reading frame (ORF) coding one of the positive regulators of σ^B^ (*rsbS*, *rsbT* and *rsbU*) or in *sigB* itself (Table 1). Additionally, one isolate (designated variant 77) harboured a 32 bp deletion in *rsbU*. As previously observed (Guerreiro, Wu, et al., 2020), all SNPs produced either frameshifts or non-sense mutations, which caused truncations of the corresponding proteins. In contrast, all grey variants isolated from the IVEE performed at 30°C carried either insertions or deletions (indels) or SNPs in the ORFs of *agrC* or *agrA*. Presumably, these mutations lead to loss-of-function of the Agr system (Table 1), resulting on the greyish colony phenotype. Additionally, none of the isolated white coloured colonies at either temperature harboured mutations in either the *sigB* or *agrCA* operons. To assess the influence of the Agr system on the colony colouration phenotype the wild type and the isogenic Δ*agrA* strains were grown separately at 30°C overnight and mixed in a ratio of 1:1 and plated in BHI agar plates at a final concentration of ~10^2^ CFU.ml^-1^. These results verified that loss of the Agr system, similarly to σ^B^, is responsible for the grey colony phenotype observed during the IVEE performed at 30°C (Fig. 1D).

**Table 1 –.**
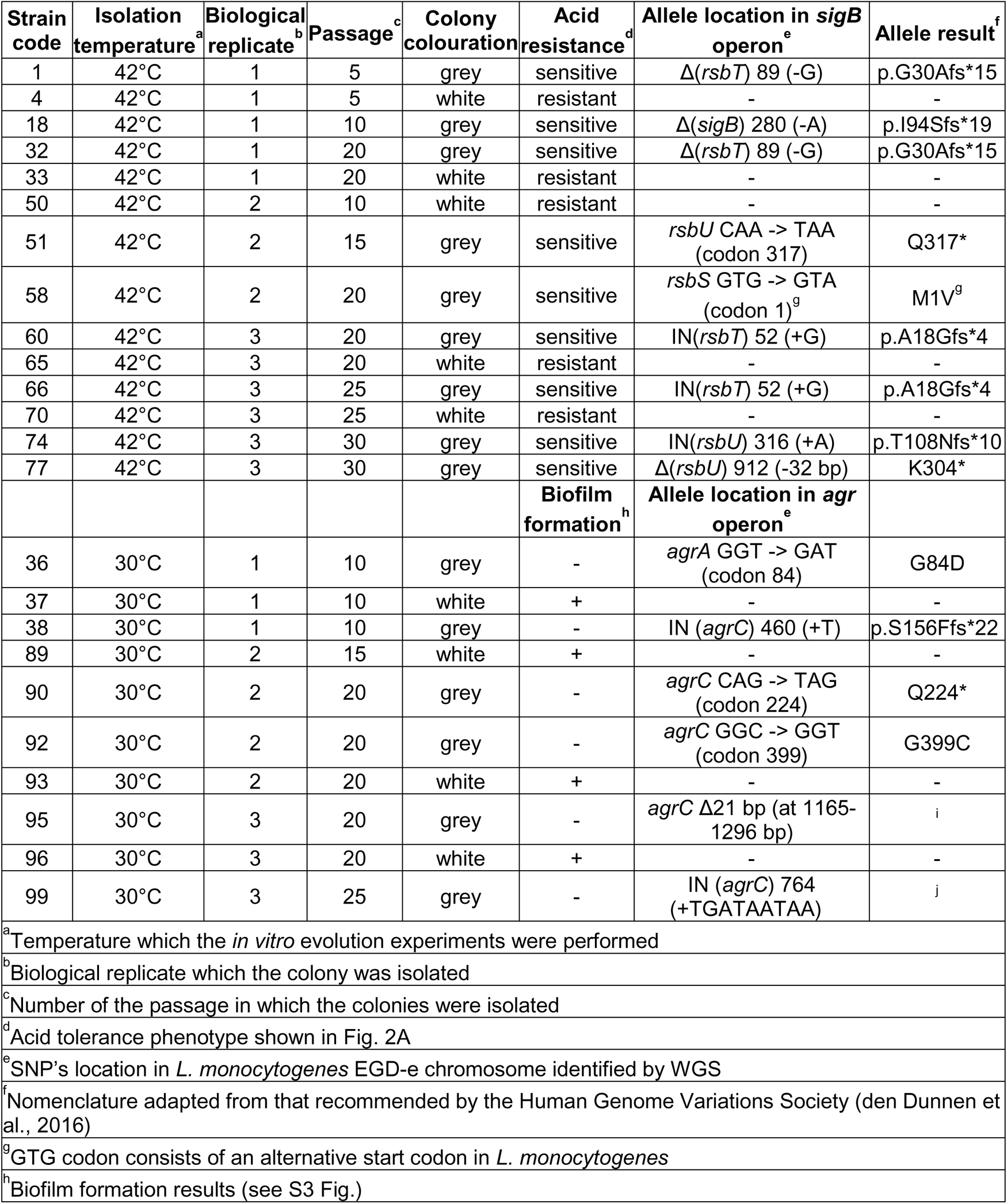
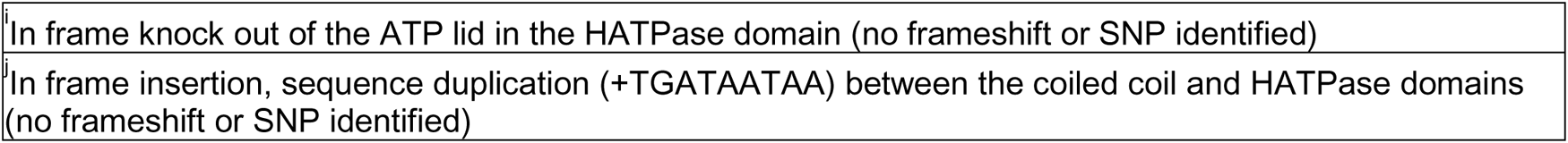
List of strains obtained during the *in intro* evolution experiments performed at 42°C and 30°C and their respective results obtained from WGS.

| Strain code | Isolation temperature<sup>a</sup> | Biological replicate<sup>b</sup> | Passage<sup>c</sup> | Colony colouration | Acid resistance<sup>d</sup> | Allele location in *sigB* operon<sup>e</sup> | Allele result<sup>f</sup> |
| --- | --- | --- | --- | --- | --- | --- | --- |
| 1 | 42°C | 1 | 5 | grey | sensitive | Δ(*rsbT*) 89 (-G) | p.G30Afs*15 |
| 4 | 42°C | 1 | 5 | white | resistant | - | - |
| 18 | 42°C | 1 | 10 | grey | sensitive | Δ(*sigB*) 280 (-A) | p.I94Sfs*19 |
| 32 | 42°C | 1 | 20 | grey | sensitive | Δ(*rsbT*) 89 (-G) | p.G30Afs*15 |
| 33 | 42°C | 1 | 20 | white | resistant | - | - |
| 50 | 42°C | 2 | 10 | white | resistant | - | - |
| 51 | 42°C | 2 | 15 | grey | sensitive | *rsbU* CAA -> TAA (codon 317) | Q317* |
| 58 | 42°C | 2 | 20 | grey | sensitive | *rsbS* GTG -> GTA (codon 1)<sup>g</sup> | M1V<sup>g</sup> |
| 60 | 42°C | 3 | 20 | grey | sensitive | IN(*rsbT*) 52 (+G) | p.A18Gfs*4 |
| 65 | 42°C | 3 | 20 | white | resistant | - | - |
| 66 | 42°C | 3 | 25 | grey | sensitive | IN(*rsbT*) 52 (+G) | p.A18Gfs*4 |
| 70 | 42°C | 3 | 25 | white | resistant | - | - |
| 74 | 42°C | 3 | 30 | grey | sensitive | IN(*rsbU*) 316 (+A) | p.T108Nfs*10 |
| 77 | 42°C | 3 | 30 | grey | sensitive | Δ(*rsbU*) 912 (-32 bp) | K304* |
|  |  |  |  |  | **Biofilm formation<sup>h</sup>** | **Allele location in *agr* operon<sup>e</sup>** |  |
| 36 | 30°C | 1 | 10 | grey | - | *agrA* GGT -> GAT (codon 84) | G84D |
| 37 | 30°C | 1 | 10 | white | + | - | - |
| 38 | 30°C | 1 | 10 | grey | - | IN (*agrC*) 460 (+T) | p.S156Ffs*22 |
| 89 | 30°C | 2 | 15 | white | + | - | - |
| 90 | 30°C | 2 | 20 | grey | - | *agrC* CAG -> TAG (codon 224) | Q224* |
| 92 | 30°C | 2 | 20 | grey | - | *agrC* GGC -> GGT (codon 399) | G399C |
| 93 | 30°C | 2 | 20 | white | + | - | - |
| 95 | 30°C | 3 | 20 | grey | - | *agrC* Δ21 bp (at 1165-1296 bp) | i |
| 96 | 30°C | 3 | 20 | white | + | - | - |
| 99 | 30°C | 3 | 25 | grey | - | IN (*agrC*) 764 (+TGATAATAA) | j |

<sup>a</sup>Temperature which the *in vitro* evolution experiments were performed
<sup>b</sup>Biological replicate which the colony was isolated
<sup>c</sup>Number of the passage in which the colonies were isolated
<sup>d</sup>Acid tolerance phenotype shown in Fig. 2A
<sup>e</sup>SNP's location in *L. monocytogenes* EGD-e chromosome identified by WGS

|  |
| --- |
| In frame knock out of the ATP lid in the HATPase domain (no frameshift or SNP identified) |
| In frame insertion, sequence duplication (+TGATAATAA) between the coiled coil and HATPase domains (no frameshift or SNP identified) |

### Acid sensitive grey coloured colonies were only isolated at 42°C

In our previous study, we reported that the loss of σ^B^ positive regulators genes (*rsbS*, *rsbU* and *rsbV*) results in an increase in acid sensitivity (Guerreiro, Wu, et al., 2020). Several of the grey coloured colonies obtained during the IVEE were challenged under lethal acidic conditions (BHI acidified to pH 2.5 at 37°C), along with a few selected white coloured isolated during the IVEE. The parental strain and white colonies exhibited similar acid tolerance, reflected by a similar pattern of survival after 30 min in pH 2.5 at 37°C (Fig. 2A). In contrast, the Δ*sigB* mutant strain and grey coloured colonies isolated at 42°C showed approximately 3 Log_10_ (CFU/ml) reduction after 30 min at pH 2.5 (Fig. 2A). At the same time, the Δ*agrA* mutant strain exhibited a small, although not significant (0.4705 Log_10_; *p* = 0.3445), reduction in resistance, while none of the grey colonies isolated at 30°C showed a reduction in acid resistance comparable to the Δ*sigB* mutant (Fig. 2B). The results suggest grey variants isolated at 42°C harbour σ^B^ loss-of-function alleles, while those isolated at 30°C do not.

**Figure 2 –.**
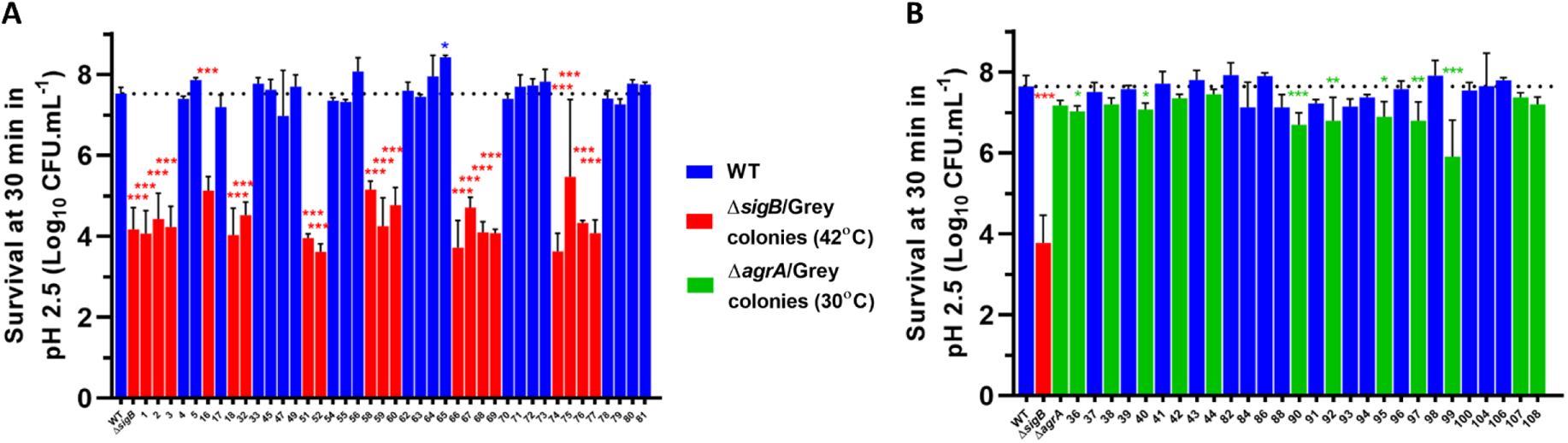
Grey colonies isolated from *in vitro* evolution experiments at 42°C have acid-sensitive phenotypes. Cultures of *L. monocytogenes* EGD-e wild-type, Δ*sigB*, Δ*agrA* and white and grey colonies isolated from the *in vitro* evolution experiments at (A) 42°C and (B) 30°C were grown overnight in BHI at 37°C. Cultures were challenged in acidified BHI medium (pH 2.5) at 37°C. Samples were taken at 0 and 30 min for viable counts. Blue bars represent white colonies strains while red bars represent the Δ*sigB* or grey colonies isolated at 42°C and green bar represents the Δ*agrA* strain or grey colonies isolated at 30°C. Two independent biological replicates were made. Statistical analysis was performed using a paired student *t*-test by comparing with the parental strain (*, *p*-value of <0.05; **, *p-*value of <0.01; ***, *p-*value of <0.001).

### Confirmation of loss-of-function alleles in the *sigB* operon and *agrCA* ORFs

The impact of the identified alleles on the activity of σ^B^ was assessed by transforming several colonies with vector pKSV7-P*_lmo2230_*::*egfp*, which harbours a σ^B^-dependent reporter (Utratna et al., 2012). The σ^B^ activity was determined through the quantification of eGFP expression in the reporter strains. Reduced σ^B^ activity was confirmed in grey coloured colonies isolated at 42°C, which harbour *sigB* alleles (Fig. 3A), while the grey colonies isolated at 30°C, harbouring alleles at the *agrCA* genes, retained similar σ^B^ activity compared with the parental strain (Fig. 3A).

**Figure 3 –.**
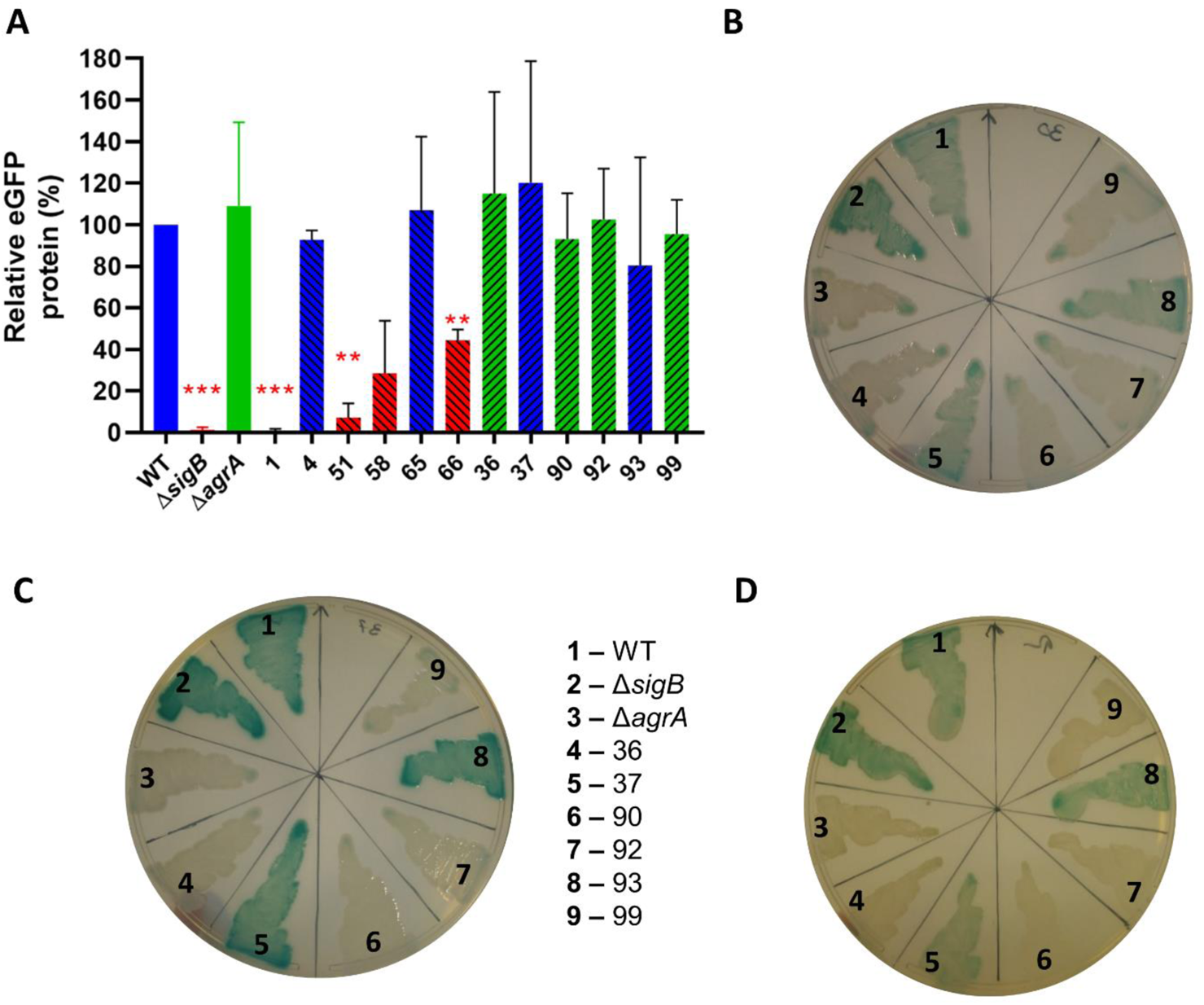
Strains carrying in alleles in *sigB* or *agr* operons exhibit reduced activity in each of the respective systems. Stationary-phase cultures of WT, Δ*sigB*, Δ*agrA* and colonies isolated from the *in vitro* evolution experiments at both temperatures, were transformed with (A) pKSV7-P*_lmo2230_*::*egfp* and (B, C, D) pGID310-P*_agrB_*::*gfp*::*bgaB*. (A) Percentage of eGFP signal normalized relative to the WT. Quantification was obtained from western blot images. Cultures transformed with pGID310-P*_agrB_*::*gfp*::*bgaB* were patched in BHI agar plates, supplemented with Chl and X-Gal, and incubated at (B) 30°C, (C) 37°C and (D) 42°C for 48 hours. For strains details refer to Table 1. Western blot quantification was generated from two independent biological replicates. Statistical analysis was performed by a paired student *t*-test (**, *p-*value of <0.01; ***, *p-*value of <0.001).

To evaluate the Agr activity, the newly constructed Agr reporter vector, pGID310-P*_agrB_*::*gfp*::*bgaB*, was transformed and integrated upstream of the original promoter region of *agrB*. Strains transformed with this reporter plasmid and with an intact Agr system form blue colonies in BHI agar supplemented with X-Gal. Reporter strains carrying *agrCA* and Δ*agrA* formed white colonies, in contrast to the blue colonies formed by the parental strain (Fig. 3B-D). Interestingly, the isogenic Δ*sigB* exhibited a slightly bluer colouration in comparison with that of the parental at all temperatures. In addition, a slight increase of *agrB* transcripts (0.95 log_2_; *p* = 0.015), a highly Agr-dependent gene, was observed in the Δ*sigB* strain (Fig. S1). These results suggest that σ^B^ exerts a repressive effect on the Agr system, an effect similar to that observed in *S. aureus* (Lauderdale et al., 2009). Finally, we observed that the activity of both reporter systems was unchanged in all the tested white colonies.

### *agr*^−^ strains exhibit a small competitive advantage against its parental strain

In our previous study, we found that *sigB*^−^ strains, exhibit increased competitive advantage at 42°C but not at 30°C when competing with the parental strain (Guerreiro, Wu, et al., 2020). To assess the competitive fitness conferred by the *sigB*^−^ alleles, competition experiments were carried out at either 30°C or 42°C using the parental strain, Δ*sigB*, and several *sigB*^−^ and *sigB*^+^ strains isolated during the IVEE at 42°C. Pairs of strains were mixed at the initial ratios of 1:1. We verified that *sigB*^−^ strains, exhibit a similar advantage over the parental and *sigB*^+^ strains at 42°C (Fig. 4A&B; Fig. S2 A&B), and as expected this phenotype was abolished at 30°C (Fig. 4C&D; Fig. S2 C&D), in agreement with our previous observations (Guerreiro, Wu, et al., 2020). The impact of the *agrCA* alleles was also assessed at 30°C by competing the parental strain against the isogenic Δ*agrA*. The results revealed a small competitive advantage that allowed the Δ*agrA* strain to reach 71.77% (*p* = 0.0066) of the total population after five passages at 30°C (Fig. 4E). Competitions between the obtained *agr*^−^ strains (colonies 36 and 92) were tested against their *agr*^+^ counterparts (colonies 37 and 93). A comparatively small competitive advantage in the *agr*^−^ strains relative to the *agr*^+^ strains was observed after 5 passages (65.7% and 61.12%; Fig. 4 F and G, respectively). However, the *agr*^−^ 36 strain could not outcompete the parental strain (Fig. S2E), while the *agr*^+^ 37 strain was overtaken by the Δ*agrA* mutant strain (Fig. S2F).

**Figure 4 –.**
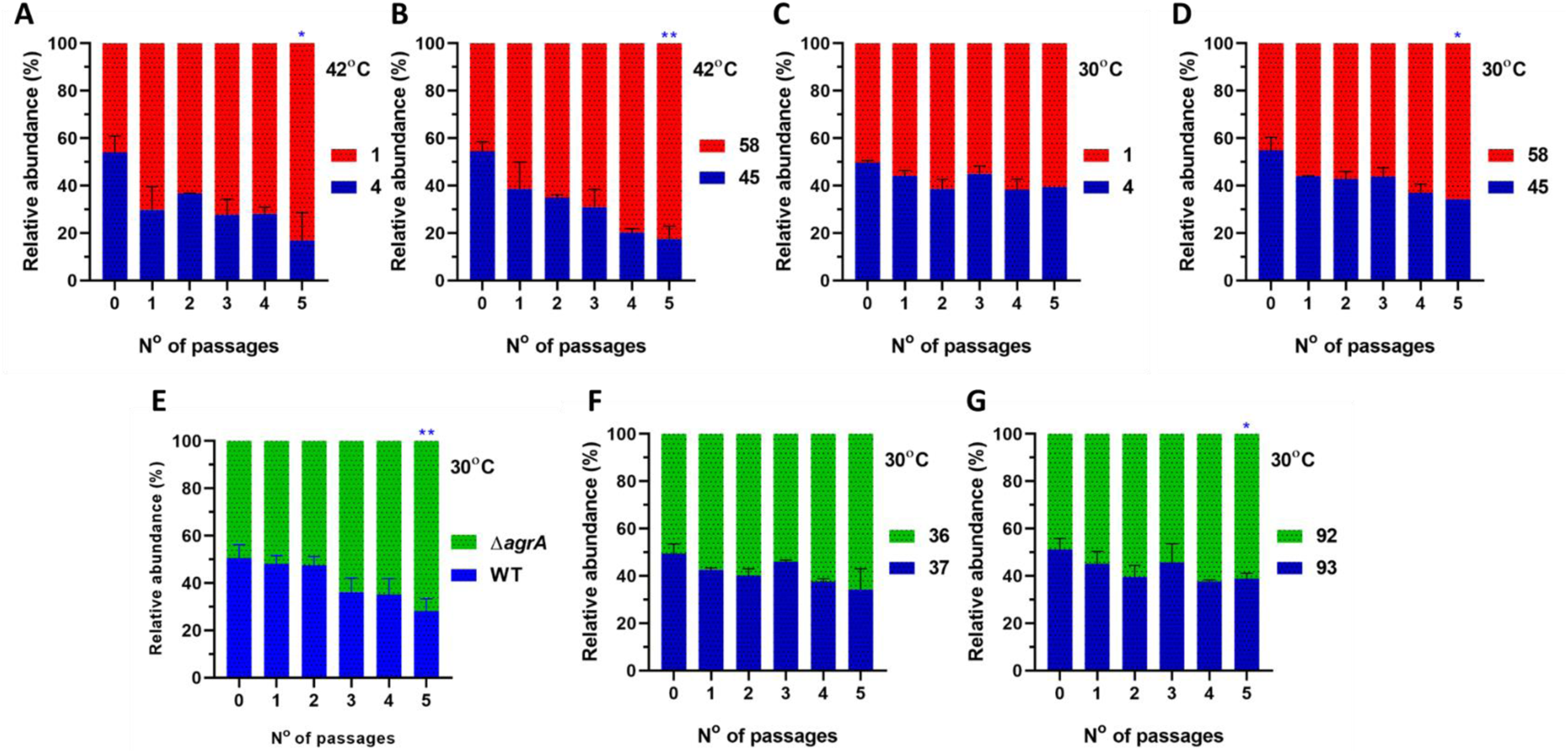
Strains carrying *sigB* or *agr* alleles exhibit a competitive advantage during mixed culture passaging. Competition experiments were performed with white versus grey colonies (starting ratio 1:1) isolated during the *in vitro* evolution experiments. Mixed cultures were passaged at (A, B) 42°C or (C, D, E, F, G) 30°C. Blue bars represent parental strain and white coloured colonies, while red and green represent *sigB*^−^ and *agr*^−^, respectively. Data were generated from two independent biological replicates. For strains details refer to Table S1. Statistical analysis was performed using a paired student *t*-test (*, *p-*value of <0.05; **, *p-*value of <0.01).

We also aimed to identify the origin of this competitive behaviour conferred by the *agrCA* alleles by measuring the growth rates of several *agr*^+^ and *agr*^−^ isolates at 30°C. However, the results showed no observable growth changes under these conditions (Fig. S2G&H). In *L. monocytogenes*, Agr activity has been linked with cell surface attachment and biofilm formation (Gray et al., 2021; Riedel et al., 2009; Rieu et al., 2007, 2008), therefore, we further tested the impact of the Agr loss-of-function alleles on the biofilm formation. The results revealed a significant reduction in the biofilm formation in Δ*agrA* and *agr*^−^ strains compared with the parental and *agr*^+^ strains (Fig. S3).

### Δ*sigB* and Δ*agrA* deletions influence the colony architecture

To our knowledge, there is no information available regarding the nature of the grey colony phenotype presented in our previous study (Guerreiro, Wu, et al., 2020). We hypothesize that such colouration differences between the parental strain and the remaining variants are a direct result of differences in the opacity of their colonies (i.e., colonies formed by the wild type were more opaque compared to those formed by *sigB^−^* and *agr^−^*). Here, 7-days-old colonies of *L. monocytogenes* EGD-e wild type, Δ*sigB* and Δ*agrA* were prepared and observed through SEM. The SEM images revealed irregular structures formed by cell aggregates at the borders of the wild type colonies (Fig. 5A), while the Δ*sigB* and Δ*agrA* mutant strains exhibited much less pronounced structures (Fig. 5B-C). Similar irregular structures were also observed while using different incubation conditions in a different study (Kumar et al., 2009). At the centre of the colonies, however, the wild type strain displayed discernible cells with a well-defined rod-like shape (Fig. 5D), whilst cells of both Δ*sigB* and Δ*agrA* mutant strains were embedded within an extracellular matrix (Fig. 5E, F). Additionally, when grown in BHI supplemented with Congo Red, both mutant strains exhibited a redder colouration compared to the wild type strain (Fig. 5G). Altogether, these data suggest that in the absence of either σ^B^ or Agr systems the colony architecture is altered following prolonged incubation on BHI agar plates, and this likely alters the opacity of the colonies resulting in a greyer appearance.

**Figure 5 –.**
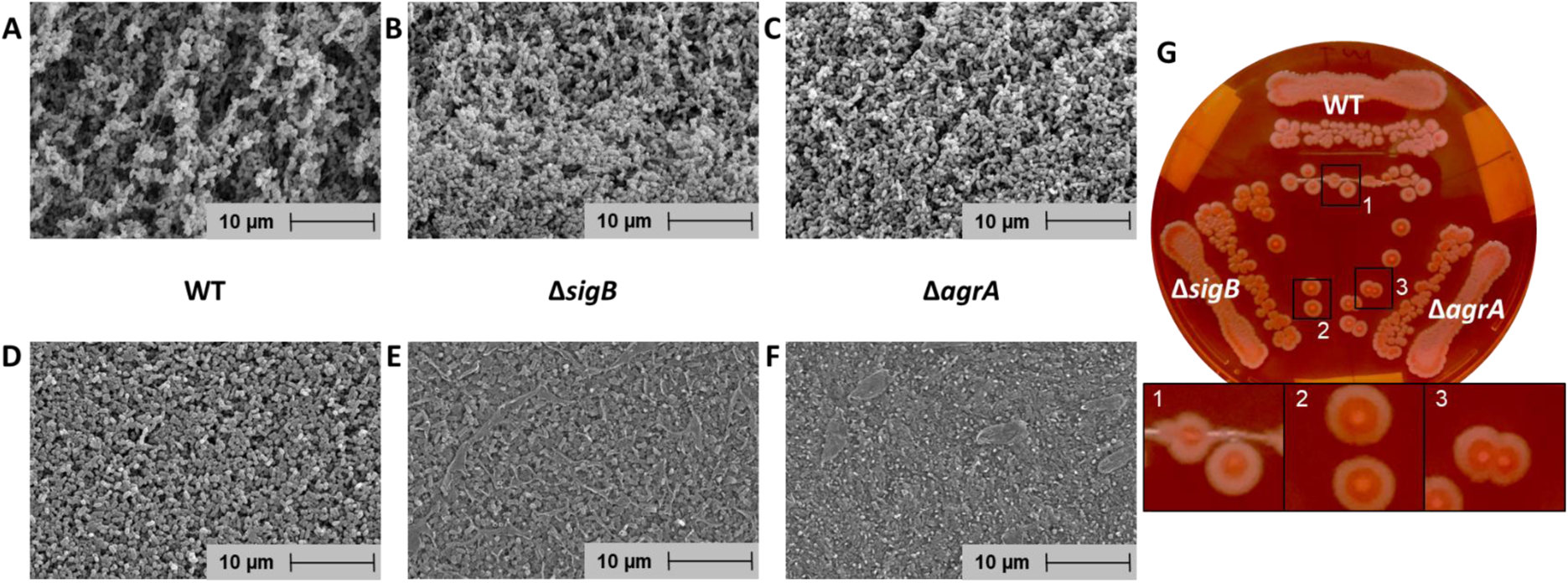
Colony architecture is influenced by either Δ*sigB* or Δ*agrA* deletions. Images recorded by scanning electron microscopy of 7-days-old colonies of (A, D) wild type, (B, E) Δ*sigB* and (C, F) Δ*agrA* grown in BHI agar plates at 30°C. Under these conditions, the grey colony phenotype was evident in the mutant strains. Images correspond to the colony (A, B, C) border or (D, E, F) centre. (G) Colonies of *L. monocytogenes* EGD-e WT, Δ*sigB* and Δ*agrA* strains grown in BHI agar supplemented with Congo Red (25 µg.ml^-1^). Plates of Congo Red were incubated for 24 h at 37°C and three more days at 30°C. Representative data from three biological replicates are shown.

### *agrCA* alleles show high PMSC rates similar to the positive regulators of σ^B^

The spontaneous emergence of *agrCA* alleles in the IVEE performed at 30°C prompted us to investigate if *agrCA* alleles also occur among wild isolates strains of *L. monocytogenes*. For this, we employed the previously described approach for analysing premature stop codon (PMSC) rates (Guerreiro, Wu, et al., 2020) to survey40,080 *L. monocytogenes* genomes publicly available on the NCBI database. We examined the PMSC occurrence rate (per 100bp) in the *agr* operon compared to genes from several other gene categories including, two-component systems, the *sigB* operon, genes used for MLST analysis, metabolic genes and virulence genes (Fig. 6). As expected, results for the *sigB* operon and other genes used as controls (MLST genes, sigma factors & metabolic genes, *rsbR1* paralogues, virulence genes) were similar to our earlier observations. The PMSC rate for *agrC* was comparable to the highest observed in *sigB* operon and was substantially higher in the case of *agrA* (Fig. 6). In clear contrast, PMSCs in all other two-component systems encoding genes were very rare, with *resD* as the sole exception. ResD is known important for mediating carbohydrate-based repression on virulence genes as well as controlling respiration in *L. monocytogenes* (Larsen et al., 2006). More interestingly, the high PMSC rates were restricted to *agrCA*, in contrast with the extremely low rates of *agrBD*. Taken together, these results indicate a high incidence of *agrCA* loss-of-function alleles among field isolates, suggesting that a selective advantage may be associated with the inactivation of AgrC and AgrA in some environments.

**Figure 6 –.**
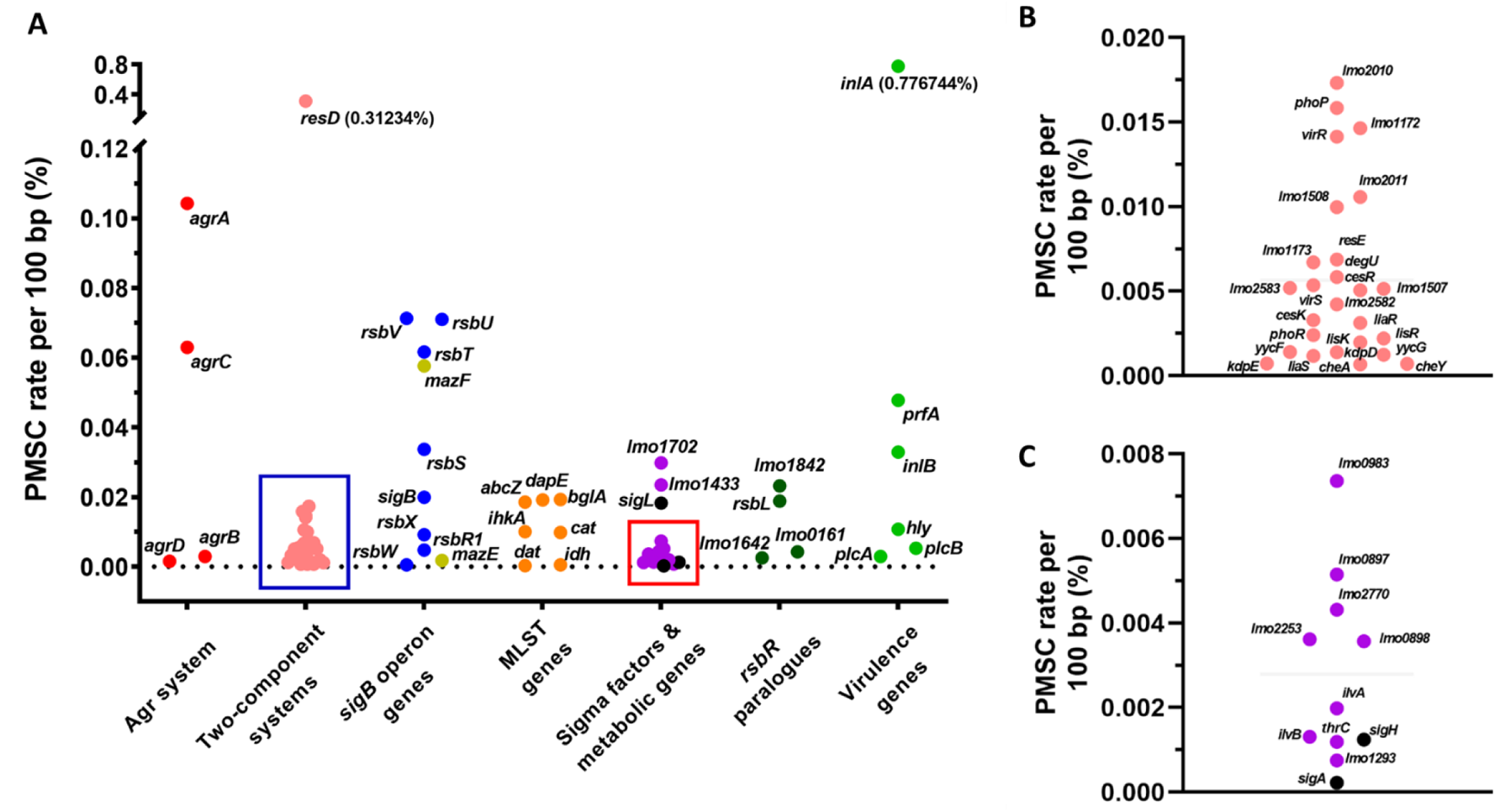
High occurrence of PMSCs in ORFs of *agrCA* and positive regulators of σ^B^. *in silico* analysis of 40,080 genomes of *L. monocytogenes* strains publicly available. (A, B and C) PMSC rate normalized by 100 bp of the open reading frames length for genes comprising the *agr* operon (red), two-component systems (pink), *sigB* operon (blue), *mazEF* (yellow), MLST genes (orange), sigma factors (black) and metabolic genes (purple), rsbR1 paralogues (dark green) and virulence genes (light green). (B and C) Expanded area indicated by the blue and red squares in panel A, respectively.

### Measurements of the Agr activity in the strains collections

Since the *in silico* analysis revealed the prevalence PMSC mutations affecting *agrC* and *agrA* AgrC and AgrA, we sought to investigate whether such mutations can influence the activity of the Agr system. An existing laboratory strain collection of 168 *L. monocytogenes* strains with WGS available (Wu & O’Byrne, unpublished data), was screened for *agr* operon mutations giving rise to PMSCs. Five strains in this collection were found to carry PMSCs in the *agrCA* genes (Table 2). The well-studied laboratory reference 10403S harbours an *agrA* mutation predicted to truncate AgrA. The clinical isolate MQ140031 bears an in-frame deletion of 141 amino acids in *agrA*. Strains 1389, 1384, and 1378 carry *agrC* alleles and are among twenty-five strains recommended for assessing food challenge studies by the European Union Reference Laboratory of *L. monocytogenes*. A truncation in the 5’-end region of *agrC* rendered the majority of the protein untranslated in strain 1389; a transposon sequence insertion in the 3’-end region of *agrC* disrupted its translation in strain 1384; a frameshift at the stop codon resulted in a longer version of *agrC* that overlaps *agrA* open reading frame in strain 1378. As these strains were from different genetic backgrounds, a control strain from the same Clonal Complex (CC) was selected for each to assess the impact of these mutations on the Agr activity. For this, eight strains were transformed and with pGID310-P*_agrB_*::*gfp*::*bgaB* and the reporter plasmid was integrated into their genomes. Agr activities were visualised on medium containing X-gal and Agr activity was determined by RT-qPCR analysis of the transcription of *agrB* and *agrA*. In both assays, the Agr activity was reduced in all strains carrying either *agrC* or *agrA* alleles (Fig. 7A&B). In conjunction with the PMSC analysis, our results suggest that, like the σ^B^ system, Agr quorum sensing system is subject to negative selective pressure under some commonly encountered conditions.

**Figure 7 –.**
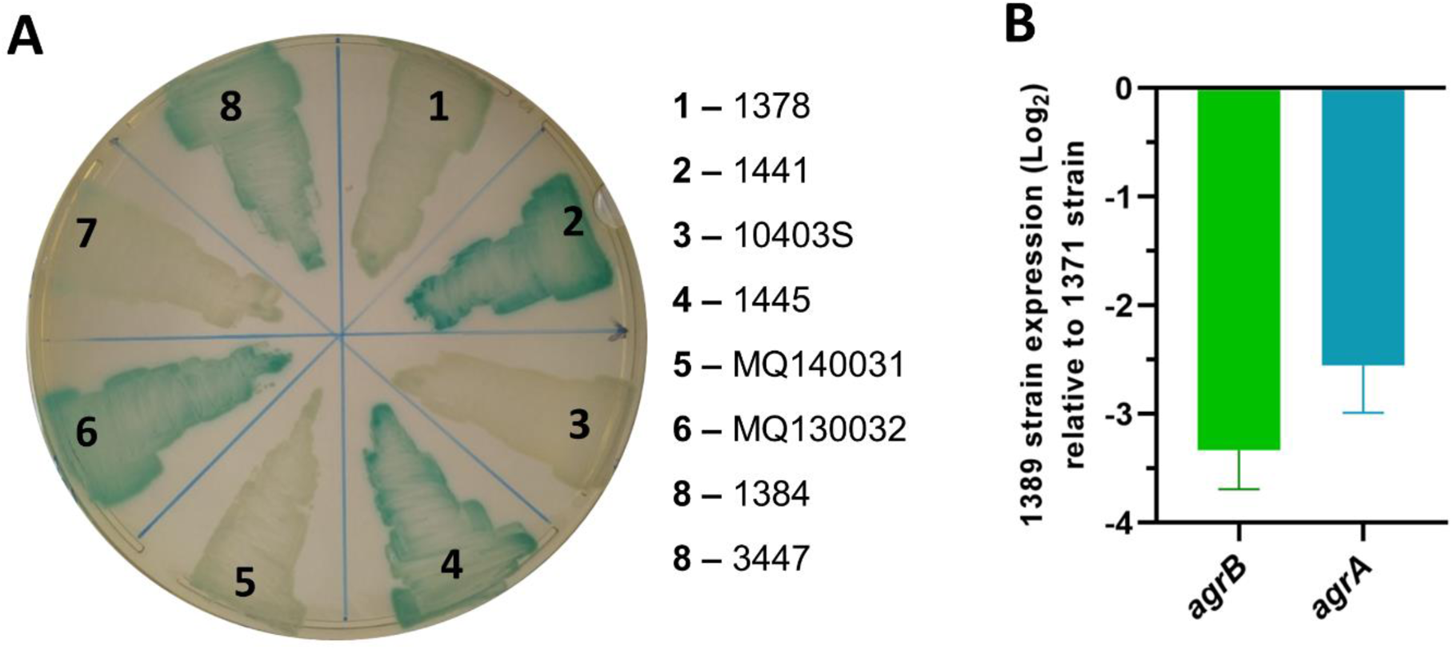
Field isolates carrying in alleles in *agr* operon exhibit reduced activity in each of the respective systems. (A) Isolates that integrated pGID310-P*_agrB_*_::_*_gfp_*_::_*_bgaB_* were patched in BHI agar plates supplemented with Chl and X-Gal and incubated at 37°C for 48 hours. (B) Expression of *agrB* and *agrA* in strain 1389 grown to stationary phase (16h) at 37°C was compared to strain 1371 by RT-qPCR experiment.

**Table 2 –.**
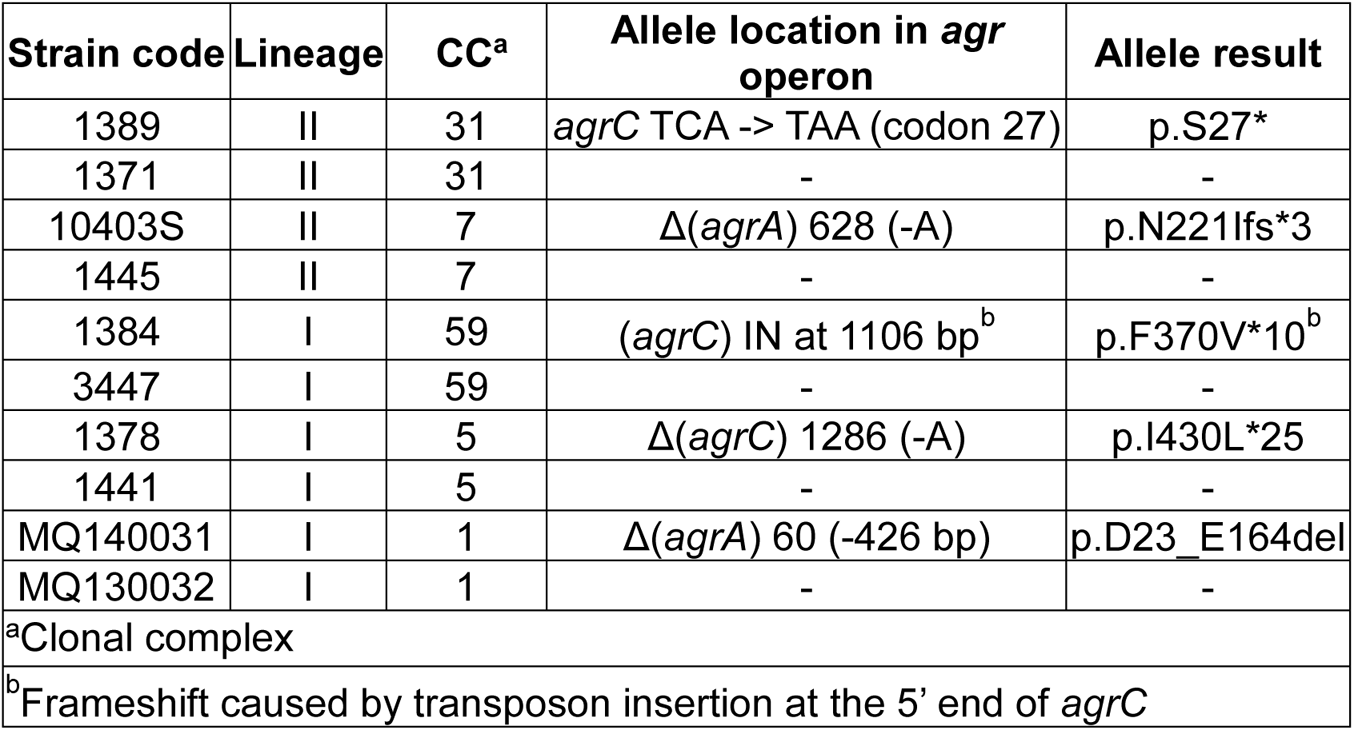
List of *L. monocytogenes* isolates that carry PMSC in *agr* operon genes and their respective reference strains that carry intact *agr* operon genes.

## Discussion

In this study, we investigated the conditions leading to the emergence and proliferation of variants harbouring alleles within the *sigB* operon in *L. monocytogenes*. We set up IVEEs at either 42°C or 30°C and, variants forming greyish more translucent colonies were detected at both temperatures (Fig. 1A-B). Grey variants isolated from the IVEE performed at 42°C harboured loss-of-function alleles at the *sigB* operon (Table 1) and exhibited reduced acid tolerance and decreased σ^B^ activity (Fig. 2A, 3A). Whereas grey variants isolated from the IVEE performed at 30°C carried loss-of-function alleles in *agrCA* genes (Table 1; Fig. 3B-D). In both cases, grey variants were able to outcompete the parental strain, at the respective temperatures, although this effect was more apparent for the *sigB* operon mutants (Fig. 4A-G). Additionally, we analysed all publicly available genomes of *L. monocytogenes* and identified high rates of PMSC occurrence in both *sigB* operon and *agrCA* ORFs (Fig. 6A).

In several other studies, loss-of-function alleles within the *sigB* operon were identified in newly constructed mutants strains or strains isolated from intragastric inoculated mice (Asakura et al., 2012; Guerreiro, Wu, et al., 2020; O’Donoghue, 2016; Quereda et al., 2013), suggesting that such alleles can emerge spontaneously in laboratory grown cultures. Indeed, we observed the spontaneous emergence of strains harbouring *sigB* alleles in our IVEE at 42°C, however, the parental strain still accounted for ~80% of the population at the end of the experiment (Fig. 1A). It is known that alleles conferring increased fitness can coexist with the parental strain for up to ~10,000 generations, until additional mutations further increase the mutant’s fitness (reviewed in (Connolly et al., 2021). However, only ~200 generations occurred during the 30 day span of our experiment. We hypothesise that *sigB*^−^ strains still require additional fitness conferring mutations before being able to fully overtake the parental strain population. While 42°C conferred increased competitive fitness to *sigB*^−^ strains (Fig. 4A&B; Fig. S2A&B; (Guerreiro, Wu, et al., 2020)), this advantage was abolished at 30°C (Fig. 4C&D; Fig. S2C&D). These observations probably explain the apparent absence of *sigB*^−^ strains in the IVEE performed at 30°C (Table 1). It is known that σ^B^ is triggered during temperature upshift (to 45-48°C) in *L. monocytogenes* (Becker et al., 1998; Guldimann et al., 2017). Moreover, under mild stresses, the σ^B^ activity is known to hinder the growth rate of *L. monocytogenes* (Brøndsted et al., 2003; Guerreiro, Wu, et al., 2020). We hypothesise that the competitive disadvantage, observed at 42°C for the wild type is a consequence of a higher activation of σ^B^, resulting in a slower growth rate. Interestingly, similar mutations in the gene encoding the alternative sigma factor σ^S^ (*rpoS*), an homologue of σ^B^ in gram-negative bacteria, are frequently found in laboratory grown *E. coli* and *Pseudomonas aeruginosa* (Robinson et al., 2020; Spira et al., 2011).

The exact mechanism responsible for the growth suppression generated by the activation of σ^B^ is still unclear, but a few hypotheses have been suggested (Guerreiro, Arcari, et al., 2020; Guerreiro, Wu, et al., 2020). First, competition between the housekeeping sigma factors A (σ^A^) and σ^B^ for the RNA polymerase may direct transcription away from genes with growth-related functions. Similarly an increase in mRNA concentration from genes belonging to the general stress response regulon will require a significant fraction of the cellular ribosome pool, thereby limiting the potential to translate housekeeping mRNA. Secondly, redirection of resources and energy from growth to survival mechanisms might negatively impact growth. Thirdly, it is possible that deliberate suppression of growth through a σ^B^-dependent mechanism might occur, as has been suggested for the σ^B^-dependent sRNA Rli47 (Marinho et al., 2019). In this scenario the biological rationale is that reduced growth might increase the likelihood of survival since it allows time for homeostatic and repair mechanisms to function. Finally, if σ^B^ plays a role in producing secreted enzymes or molecules that can be shared within the population, mutants could arise as cheaters within the population that take advantage by exploiting the availability “public” resources. Further experimental work will be needed to investigate which of these explanations is driving the selective pressure for σ^B^ loss-of-function mutants to emerge at 42°C.

The IVEE performed at 30°C revealed grey variants harbouring alleles in the ORF of either *agrC* or *agrA* instead of PMSC in the *sigB* operon (Fig.1; Table 1), which lead to the inactivation of the Agr system (Fig. 3B-D). Additionally, the *agr*^−^ alleles seem to provide a small competitive fitness advantage at 30°C (Fig. 4E-G), however, the growth rates of these variants remained unchanged (Fig. S2G & H). Interestingly, the *agr*^−^ alleles impaired the biofilm formation in conical-bottomed 50ml plastic tubes under shaking conditions (Fig. S3). The Agr system plays a critical role in the surface cell attachment during the initial stages of the biofilm formation (Rieu et al., 2008). We speculate that a small fraction of the parental strain cells attach and form biofilm on the wall of the flask, while the emerging *agr*^−^ cells, unable to attach to the flasks surfaces, are left in the planktonic phase. This phenotype could contribute to the a disproportional ratio of parental:*agr*^−^ that is magnified with the successive passages. Interestingly, while the Agr activity is crucial for the biofilm formation at low temperatures, this phenotype is reverted at 37°C (Garmyn et al., 2012), which may explain the absence of Agr alleles detected in the IVEE performed at 42°C. Other non-exclusive factors may contribute to the emergence of *agr* mutations, as the Agr and other QS systems represent an energetic burden and detrimental for growth in certain environments (García-Contreras et al., 2015; George et al., 2019; Whiteley et al., 2017). In a recent study, the *L. monocytogenes* Agr system was found to downregulate genes involved in the synthesis of S-adenosylmethionine in nutrient-poor media (Garmyn et al., 2012; Lee & Wang, 2020), suggesting that Agr represses metabolism under specific conditions.

The high frequencies of PMSC in *agrCA* genes amongst publically available *L. monocytogenes* genome sequences contrast with an extremely low frequency of PMSC found in *agrBD* genes (Fig. 6). In accordance with this, only *agrCA* alleles emerged during the *in vitro* evolution experiment at 30°C. The *agrCA*^−^ strains may behave as social cheaters, which still contribute to the extracellular auto-inducing peptide (AIP) pool and stimulate the Agr activity on *agr*^+^ strains. However, these strains do not bear the energetic burden of contributing to the community. To our knowledge, the contribution of the Agr system in the production of public goods in *L. monocytogenes* communities is currently uncharacterized. Our data suggest that similarly to σ^B^, the deployment of the Agr system possesses a fitness trade-off for *L. monocytogenes* under specific conditions.

In our previous study, we found that Δ*sigB* and *sigB*^−^ strains formed greyish and more translucent colonies in comparison with the parental strain (Guerreiro, Wu, et al., 2020). Here, we found that Δ*agrA* and *agr*^−^ strains form similar colonies to those of *sigB*^−^ strains (Fig. 1 D). Previous studies found that σ^B^ is responsible for the cell wall thickness in response to blue light stress, which in turn impacts the colony opaqueness (O’Donoghue et al., 2016; Tiensuu et al., 2013). Interestingly, the Agr system also regulates the expression of proteins participating in the cell wall peptidoglycan biosynthesis in *L. monocytogenes* (Lee & Wang, 2020). It is plausible that the colouration disparities observed in this study are a consequence of differences in cell envelope composition, however, future studies will have to confirm this hypothesis.

SEM revealed complex multicellular structures close to the colony edges of the parental colonies (Fig. 5A), while cells at the centre were compact and well defined (Fig. 5D). These structures were less evident in both Δ*sigB* and Δ*agrA* strains (Fig. 5B-C), while their colony centres apparently had a more abundant extracellular matrix (Fig. 5E-F), which seems correlated with the red colouration observed in plates supplemented with Congo red (Fig. 5G). *S. aureus* is known to form irregular biofilms that is dependent on the production of the Agr-dependent phenol-soluble modulins (PSM) (Periasamy et al., 2012), however, PSM have not been identified in *L. monocytogenes*. It is not surprising that both σ^B^ and Agr regulate biofilm formation as previous studies demonstrated an overlap in their regulons (Garmyn et al., 2012; Riedel et al., 2009), however the environmental signals and genes responsible for these phenotypes are currently unknown.

In summary, our results further suggest that laboratory conditions could inadvertently select unwanted secondary mutations in non-essential or energetic expensive systems. These findings highlight the need for caution in the routine culturing of laboratory stocks of *L. monocytogenes* and especially during the execution of mutagenesis protocols that require an elevated temperature incubation, to avoid inadvertently picking up a mutation in one of these systems. Such mutations could easily confound any phenotypic study of this pathogen, potentially leading to erroneous conclusions. Its seems appropriate therefore to suggest that whole-genome sequencing should be implemented as a standard verification protocol for any newly constructed mutants to avoid misinterpretation of phenotypical data.

## Material and methods

### Bacterial strains, plasmids and primers

*L. monocytogenes* and *E. coli* strains used in this study are shown in Table S1. Primer sequences and plasmids are shown in Table S2. Strains were grown in brain heart infusion (BHI) broth or agar (LabM™) at 37°C with constant shaking of 150 rpm, unless otherwise specified. Cells were grown overnight for 16 hours to reach the stationary phase. Antibiotics were added to the growing media when required at the following concentrations: ampicillin (Amp) at 100 µg.ml^-1^ for the *E. coli* strain; chloramphenicol (Chl) at 10 µg.ml^-1^, tetracycline (Tet) at 2.5 µg.ml^-1^ for *L. monocytogenes* strains.

### Long-term incubation *in vitro* evolution setup

*L. monocytogenes* EGD-e wild type strain was grown overnight at either 30°C or 42°C in 50 mL conical bottom flasks containing 5 mL of fresh BHI. Each overnight culture was slipped in two flasks through a dilution of 1:100 and allowed to grow for 24 hours at the respective temperatures. Passages consisting of a 1:100 dilution of each culture into fresh BHI was made every 24 hours for 30 days. Sampling was made every five passages by serial diluting in PBS (Sigma^®^) to 10^-7^ and plating in BHI agar plates. Plates were incubated at 37°C for 24 hours and further 6 days at 30°C. Colonies colouration was registered and relative abundance was calculated as the percentage of the total number of colonies. The white/grey colony colourations differentiated the non-mutated parental from strains carrying putative alleles in the *sigB* and *agr* operons. A total of six flasks, two per biological replicate, were used at each temperature.

### Congo red agar plates

Cultures of *L. monocytogenes* WT, Δ*sigB* and Δ*agrA* were separately grown in BHI overnight at 30°C. Cultures were serially diluted in PBS (Sigma^®^) until 10^-7^ and 10 µl of the dilutions between 10^-5^ and 10^-7^ were plated in BHI agar plates supplemented with Congo Red (Sigma^®^) at a final concentration of 25 µg.ml^-1^. Plates were incubated for 24 h at 37°C and transferred to 30°C for three more days until the differentiation in colony colouration was observable.

### Acid tolerance in BHI at pH 2.5

Stationary-phase cultures of the wild type, Δ*sigB,* Δ*agrA* and mutant strains were centrifuged at 14,000 rpm for 1 min and resuspended in BHI medium previously acidified with 5 M HCl to pH 2.5. Resuspensions were incubated in a water bath at 37°C for 30 min. Samples were taken at 0 and 30 min, serially diluted from 10^-7^ to 10^-2^ in PBS (Sigma^®^) and plated onto BHI agar plates. Plates were incubated at 37°C and colonies were counted 24 hours afterwards. Two biological replicates were made.

### Whole-genome sequencing

The gDNA of strains obtained during the IVEE at both temperatures was extracted with DNeasy^®^ Blood and Tissue Kit (Qiagen) according to the manufacturer’s recommendations. Purified gDNA was sent to MicrobesNG for WGS and the retrieved trimmed reads were used for SNP analysis performed in Breseq (Deatherage & Barrick, 2014). The chromosomal sequence of *L. monocytogenes* EGD-e (NCBI RefSeq accession no. NC_003210.1) was used as the reference genome.

### Construction of the *agr* system reporter pGID310 (P*_agrB_*::*gfp*::*bgaB*) plasmid

The Agr reporter plasmid was constructed by amplifying the *bgaB* ORF from *Geobacillus stearothermophilus*, contained in pDL (Yuan & Wong, 1995), using the primers Bga11 and Bga12. The resulting amplicon and pGID128 were separately digested using the restriction enzymes BglII (ThermoFisher). The digested DNA was ligated using T4 ligase (Promega) and subsequently transformed into *E. coli* One Shot™ TOP10 (Invitrogen). The resulting plasmid was named pGID310-P*_agrB_*::*gfp*::*bgaB*.

### Bacterial transformation with σ^B^ and Agr reporter plasmids

*L. monocytogenes* electrocompetent cells were created as previously described (Monk et al., 2008). Cells were transformed with either pKSV7-P*_lmo2230_*::*egfp* (Utratna et al., 2012) or pGID310-P*_agrB_*::*gfp*::*bgaB*. Transformed colonies were selected from BHI agar plates containing Chl and incubated at 30°C. Chromosomal integration of the plasmids was carried out as achieved as previously described (Utratna et al., 2012), by incubating fluorescent cells at 42°C. The pKSV7-P*_lmo2230_*::*egfp* chromosomal integration occurred upstream of the original P*_lmo2230_* through homologous recombination and pGID310-P*_agrB_*::*gfp*::*bgaB* integrated upstream of the original P*_agrB_*. Integration was verified by colony PCR using the forward primers *egfp*-*lmo2230*-F for pKSV7-P*_lmo2230_*::*egfp* and *bgaB*_*agrB*_F for pGID310-P*_agrB_*::*gfp*::*bgaB* in conjugation with the reverse primer *egfp*/*bgaB*_*lmo2230*_R for both plasmids.

### Western blots for eGFP protein

*L. monocytogenes* EGD-e wild type, Δ*sigB*, Δ*agrA* and strains, obtained during the *in vitro* evolution experiment at 42°C and 30°C, were grown to stationary phase at 37°C and the total protein fractions were extracted. Cultures were supplemented with Tet and centrifuged at 9,000 x g for 15 min at 4°C. Pellets were resuspended in sonication buffer (13 mM Tris-HCl, 0.123 mM EDTA, and 10.67 mM MgCl2, adjusted to pH 8.0). Bacterial suspensions were digested with 1 mg.ml^-1^ of lysozyme (Sigma^®^) for 30 min and centrifuged once more and resuspended in sonication buffer. Resuspensions were then transferred into cryotubes containing zirconia-silica beads (Thistle Scientific) and bead beaten in a FastPrep-24™ at a speed of 6 m.s^-1^ for 40 s, twice. Cell lysates were centrifuged at 13,000 x g for 30 min at 4°C and the supernatant was recovered. Total protein quantification was performed using Pierce™ BCA Protein Assay Kit (Thermo Scientific) according to the manufacturer’s recommendations. Total protein concentrations were adjusted to a final concentration of 0.5 mg.ml^-1^, and 12 µL. Each sample was separated by SDS-PAGE (15% acrylamide/Bis-acrylamide), along with the PageRuler™ Plus Prestained protein ladder (Thermo Scientific). Protein was transferred onto a polyvinylidene difluoride (PVDF) membrane and blocked with Tris-Buffered Saline supplemented with 0.1% (vol/vol) of Tween 20 (Sigma^®^) (TBST) supplemented with 3% (wt/vol) skim milk powder (Sigma^®^). SDS-PAGE gels were stained with GelCode^®^ Blue Staining Reagent (Thermo Scientific) and later destained in distilled water. Immunoblots were probed with rabbit polyclonal IgG anti-GFP antibodies (Santa Cruz Biotechnology^®^) diluted 1:500 in TBST and incubated overnight at 4°C. The secondary antibody mouse anti-rabbit IgG–horseradish peroxidase (HRP) (Santa Cruz Biotechnology^®^) was diluted to a 1:6,500 ratio in TBST and incubated with the membrane at RT for 1 hour. Immunoblots were visualized in the Odyssey Fc imaging system (Li-Cor^®^ Biosciences). Band intensity was registered and quantified in Image Studio Lite ver5.2. Results are presented as a percentage of relative eGFP protein. Two biological replicates were made.

### Qualitative analysis of the *agr* activity

*L. monocytogenes* EGD-e wild type, Δ*sigB*, Δ*agrA* and strains obtained the IVEE at 30°C and transformed with pGID310 were grown overnight in BHI supplemented with Chl at 30°C and streaked in BHI agar plates supplemented with X-Gal 0.1 mg.mL^-1^ (Thermo Scientific) and incubated for at 30°C, 37°C or 42°C. White or blue colony colouration was registered after 48 hours of incubation.

### *In vitro* competition experiments

*L. monocytogenes* EGD-e wild type, Δ*sigB*, Δ*agrA* and strains obtained during the IVEE at both temperatures were grown overnight in BHI at either 30°C or 37°C. Competitions experiments were carried out as previously described (Guerreiro, Wu, et al., 2020). Briefly, stationary phase cultures were adjusted to an initial OD_600_ of 0.05 in fresh BHI medium at final ratios of 1:1 and incubated at the indicated temperatures. Passages were made every 24 hours by diluting the cultures 1:100 into fresh BHI medium for 5 days. Samples were taken in every passage, diluted to 10^-7^ in PBS, plated onto BHI plates and incubated at 37°C for 24 hours and further incubated at 30°C for 5 days. Relative abundance was calculated.

### Scanning electron microscopy

Overnight cultures of *L. monocytogenes* EGD-e wild type, Δ*sigB* and Δ*agrA* were diluted and plated in BHI agar plated at a concentration of 10^2^ CFU.mL^-1^. Plates were incubated at 37°C for 24 hours and further incubated at 30°C for 6 days. Squares of approximately 1 cm^2^ were cut from the agar containing a single colony were cut and submerged in EM fixative buffer (2% glutaraldehyde + 2% paraformaldehyde in 0.1M sodium cacodylate buffer, pH 7.2) for 2 hours at RT. EM fixative buffer was replaced by 0.2 M sodium cacodylate (pH 7.2) and stored at 4°C until further processed. Samples were dehydrated for 15 min in successive immersions in increasing concentrations of ethanol (30%, 50%, 70%, 90% and 100%), twice per concentration. Samples were then dried by Critical Point Drying (Leica EM CPD300) and sputter-coated with gold (Quorum Q150R ES Plus). Samples were visualized in SEM (Hitachi S2600N). Three biological replicates were made.

### *In silico* analysis of premature stop codon occurrence rates

Methodology for this analysis was directly adapted from our previous study (Guerreiro, Wu, et al., 2020) however, with 40,080 genome sequences available from the NCBI database analysed (accessed April 19^th^ 2021). Briefly, a BLAST DNA database was constructed for each genome and Blastn match for each gene of interest was extracted and translated *in silico*. For the genes determined as present in the genome, lengths of deduced translation products were calculated as percentage values relative to the length of reference protein sequence from *L. monocytogenes* EGD-e strain. PMSC is defined as <90% of reference protein length and its occurrence as a rate corrected for gene length. In addition to the 41 genes previously analysed, *agrBDCA* and all 31 genes encoding two-component systems, histidine kinases or response regulators are included (Williams et al., 2005). The gene *lmo1508* that encodes for a histidine kinase is among MLST genes and thus presented only in the two-component systems group (Fig. 6). Two of the two-component system encoding genes (*lmo1060* and *lmo1061*) were excluded from Figure 6 as it is absent in strains from *L. monocytogenes* lineage I.

### RT-qPCR

Cultures of *L. monocytogenes* EGD-e wild type, Δ*sigB*, Δ*agrA* and Δ*sigB*;Δ*agrA* double mutant were grown in BHI to mid-log phase (OD_600_ = 0.4) at 30°C. For analysis on field isolates, strains were grown in BHI to stationary phase at 37°C. RNAlater (Sigma^®^) was used to stop the transcription. Extraction of total RNA was made with RNeasy Minikit (Qiagen) according to the manufacturer’s recommendations. Cells lysis was achieved by bead beating twice in a FastPrep-24 at a speed of 6 m.s^-1^ for 40 s. DNA was digested with Turbo DNA-free (Invitrogen) according to the manufacturer’s recommendations. RNA integrity was verified by 0.7% agarose gel electrophoresis. SuperScript III first-strand synthesis system (Invitrogen) was used to synthesize cDNA, according to the manufacturer’s recommendations and further quantified using Qubit (Invitrogen). RT-qPCR was performed using a QuantiTect SYBR green PCR kit (Qiagen) and primers for the target genes (Table S2). Primer efficiencies for the target genes 16S, *lmo2230*, *agrA* and *agrB* were determined using purified *L. monocytogenes* gDNA. Samples were analysed on a LightCycler 480 system (Roche) with the following parameters: 95°C for 15 min; 45 cycles of 15 s at 95°C, 15 s at 53°C, and 30 s at 72°C; a melting curve drawn for 5 s at 95°C and 1 min at 55°C, followed by increases of 0.11°C.s^-1^ until 95°C was reached; and cooling for 30 s at 40°C. Cycle quantification was calculated by using LightCycler 480 software version 1.5.1 (Roche) and the Pfaffl relative expression formula (Pfaffl, 2001; Pfaffl et al., 2002). 16S rRNA expression was used as a reference. Three biological replicates were performed. Results were converted to Log_2_ expression normalized relative to the WT strain.

### Growth kinetics

Several *L. monocytogenes* strains were grown to stationary phase in BHI medium at 30°C were adjusted to an initial OD_600_ of 0.05 and grown in fresh BHI medium at 30°C. Measurements of the OD_600_ of each culture were made every hour for 8 hours. Growth rates were calculated with GrowthRates 4.3 software (Hall et al., 2014). Two biological replicates were made.

### Biofilm formation

*L. monocytogenes* strains were grown overnight in BHI at 30°C. Cultures were diluted 1:100 into 50 mL conical bottom flasks containing 5 mL fresh BHI and further incubated for 24 hours. Liquid cultures were discarded, washed twice with PBS and dried at RT for 30 min. Flasks were dyed with 1% Crystal Violet (Sigma-Aldrich^®^) for 30 min at RT and further washed twice with PBS and once with deionized water. Flasks were dried overnight at RT and pictures were taken. Two biological replicates were made.

## Acknowledgments

The authors would like to thank members of the PATHSENSE network (https://www.pathsense.eu/) for helpful discussions throughout this study. This project has received funding from the European Union’s Horizon 2020 research-and-innovation program under Marie Skłodowska-Curie grant agreement no. 721456. Jialun Wu was funded by the Department of Agriculture, Food and the Marine (17/F/244).

## Supplemental material

**Table S1 –.** List of *L. monocytogenes* strains used in this study.

| **Strains** | **Reference** |
| --- | --- |
| *Escherichia coli* One Shot™ TOP10 | Invitrogen |
| *Listeria monocytogenes* EGD-e (wild type) | K. Boor |
| *L. monocytogenes* EGD-e; $\Delta sigB$ | (1) |
| *L. monocytogenes* EGD-e; $\Delta agrA$ | (2) |
| *L. monocytogenes* 1378 |  |
| *L. monocytogenes* 1441 |  |
| *L. monocytogenes* 10403S | K. Boor |
| *L. monocytogenes* 1445 |  |
| *L. monocytogenes* MQ140031 | (3) |
| *L. monocytogenes* MQ130032 | (3) |
| *L. monocytogenes* 1384 |  |
| *L. monocytogenes* 3447 |  |
| *L. monocytogenes* 1389 |  |
| *L. monocytogenes* 1371 |  |
| **IVEE isolated strains (isolation temperature)** |  |
| *L. monocytogenes*; 1 (42°C) | This study |
| *L. monocytogenes*; 4 (42°C) | This study |
| *L. monocytogenes*; 18 (42°C) | This study |
| *L. monocytogenes*; 32 (42°C) | This study |
| *L. monocytogenes*; 33 (42°C) | This study |
| *L. monocytogenes*; 50 (42°C) | This study |
| *L. monocytogenes*; 51 (42°C) | This study |
| *L. monocytogenes*; 58 (42°C) | This study |
| *L. monocytogenes*; 60 (42°C) | This study |
| *L. monocytogenes*; 65 (42°C) | This study |
| *L. monocytogenes*; 66 (42°C) | This study |
| *L. monocytogenes*; 70 (42°C) | This study |
| *L. monocytogenes*; 74 (42°C) | This study |
| *L. monocytogenes*; 77 (42°C) | This study |
| *L. monocytogenes*; 36 (30°C) | This study |
| *L. monocytogenes*; 37 (30°C) | This study |
| *L. monocytogenes*; 38 (30°C) | This study |
| *L. monocytogenes*; 89 (30°C) | This study |
| *L. monocytogenes*; 90 (30°C) | This study |
| *L. monocytogenes*; 92 (30°C) | This study |
| *L. monocytogenes*; 93 (30°C) | This study |
| *L. monocytogenes*; 95 (30°C) | This study |
| *L. monocytogenes*; 96 (30°C) | This study |
| *L. monocytogenes*; 99 (30°C) | This study |
| **Reporter strains** |  |
| *L. monocytogenes* (wild type) + pKSV7-P<sub>Imo2230</sub>::*egfp* | (4) |
| *L. monocytogenes*; $\Delta sigB$ + pKSV7-P<sub>Imo2230</sub>::*egfp* | (1) |
| *L. monocytogenes*; $\Delta agrA$ + pKSV7-P<sub>Imo2230</sub>::*egfp* | This study |

**Table S1** – List of *L. monocytogenes* strains used in this study (continuation).
|  |  |
| --- | --- |
| *L. monocytogenes* (wild type) + pGID310-P<sub>agrB</sub>::gfp::bgaB | This study |
| *L. monocytogenes*; ΔsigB + pGID310-P<sub>agrB</sub>::gfp::bgaB | This study |
| *L. monocytogenes*; ΔagrA + pGID310-P<sub>agrB</sub>::gfp::bgaB | This study |
| *L. monocytogenes*; 1 + pKSV7-P<sub>lmo2230</sub>::egfp | This study |
| *L. monocytogenes*; 4 + pKSV7-P<sub>lmo2230</sub>::egfp | This study |
| *L. monocytogenes*; 51 + pKSV7-P<sub>lmo2230</sub>::egfp | This study |
| *L. monocytogenes*; 58 + pKSV7-P<sub>lmo2230</sub>::egfp | This study |
| *L. monocytogenes*; 65 + pKSV7-P<sub>lmo2230</sub>::egfp | This study |
| *L. monocytogenes*; 66 + pKSV7-P<sub>lmo2230</sub>::egfp | This study |
| *L. monocytogenes*; 36 + pKSV7-P<sub>lmo2230</sub>::egfp | This study |
| *L. monocytogenes*; 37 + pKSV7-P<sub>lmo2230</sub>::egfp | This study |
| *L. monocytogenes*; 90 + pKSV7-P<sub>lmo2230</sub>::egfp | This study |
| *L. monocytogenes*; 92 + pKSV7-P<sub>lmo2230</sub>::egfp | This study |
| *L. monocytogenes*; 93 + pKSV7-P<sub>lmo2230</sub>::egfp | This study |
| *L. monocytogenes*; 99 + pKSV7-P<sub>lmo2230</sub>::egfp | This study |
| *L. monocytogenes*; 36 + pGID310-P<sub>agrB</sub>::gfp::bgaB | This study |
| *L. monocytogenes*; 37 + pGID310-P<sub>agrB</sub>::gfp::bgaB | This study |
| *L. monocytogenes*; 90 + pGID310-P<sub>agrB</sub>::gfp::bgaB | This study |
| *L. monocytogenes*; 92 + pGID310-P<sub>agrB</sub>::gfp::bgaB | This study |
| *L. monocytogenes*; 93 + pGID310-P<sub>agrB</sub>::gfp::bgaB | This study |
| *L. monocytogenes*; 99 + pGID310-P<sub>agrB</sub>::gfp::bgaB | This study |
| *L. monocytogenes* 1378 + pGID310-P<sub>agrB</sub>::gfp::bgaB | This study |
| *L. monocytogenes* 1441 + pGID310-P<sub>agrB</sub>::gfp::bgaB | This study |
| *L. monocytogenes* 10403S + pGID310-P<sub>agrB</sub>::gfp::bgaB | This study |
| *L. monocytogenes* 1445 + pGID310-P<sub>agrB</sub>::gfp::bgaB | This study |
| *L. monocytogenes* MQ140031 + pGID310-P<sub>agrB</sub>::gfp::bgaB | This study |
| *L. monocytogenes* MQ130032 + pGID310-P<sub>agrB</sub>::gfp::bgaB | This study |
| *L. monocytogenes* 1384 + pGID310-P<sub>agrB</sub>::gfp::bgaB | This study |
| *L. monocytogenes* 3447 + pGID310-P<sub>agrB</sub>::gfp::bgaB | This study |

**Table S2 –.**
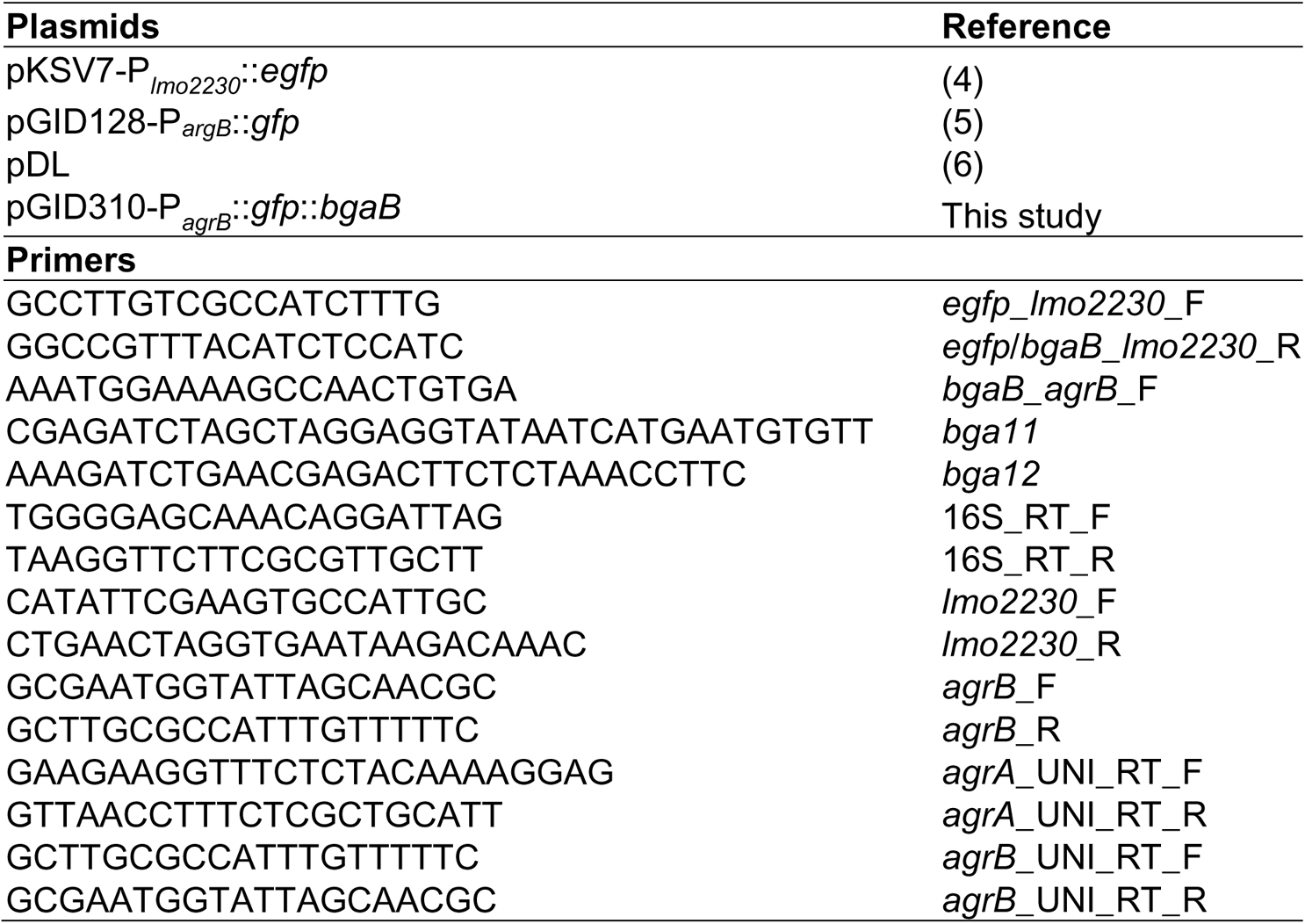
List of plasmids and primers used in this study.

**Figure S1 –.**
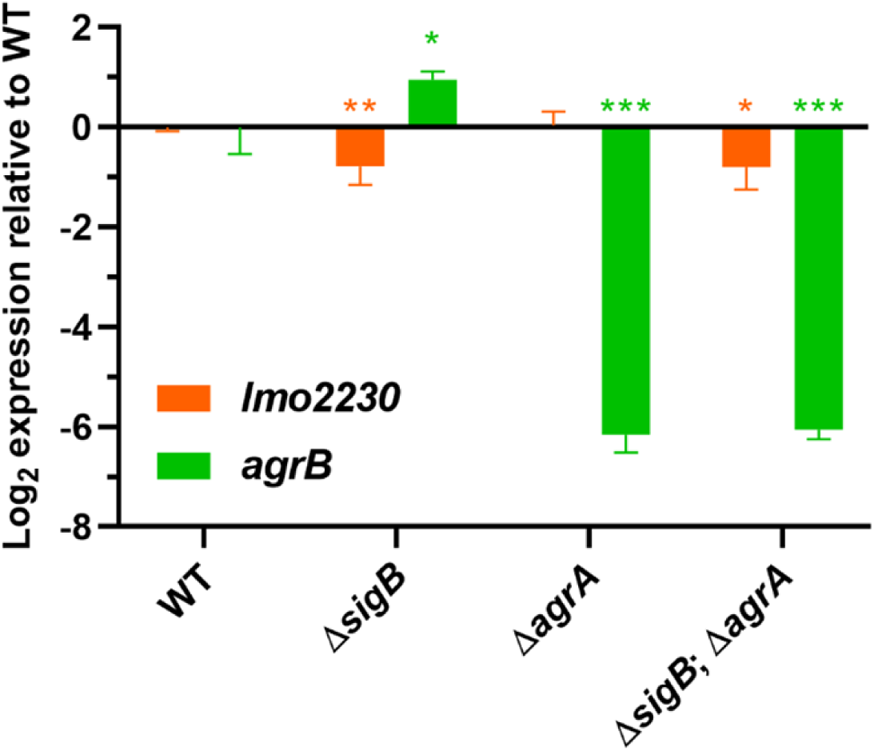
Evidence for limited cross-talk between the σ^B^ and Agr systems at 30°C. Expression of *lmo2230* and *agrB* measurements by RT-qPCR in wild type, Δ*sigB*, Δ*agrA* and the double mutant Δ*sigB* Δ*agrA* grown in BHI at 30°C. Mid-log phase cultures (OD_600_ = 0.4) were used for these results. Three independent biological replicates were made. Statistical analysis was performed using a paired student *t*-test (*, *p-*value of <0.05; **, *p-*value of <0.01; ***, *p*-value of <0.001).

**Figure S2 –.**
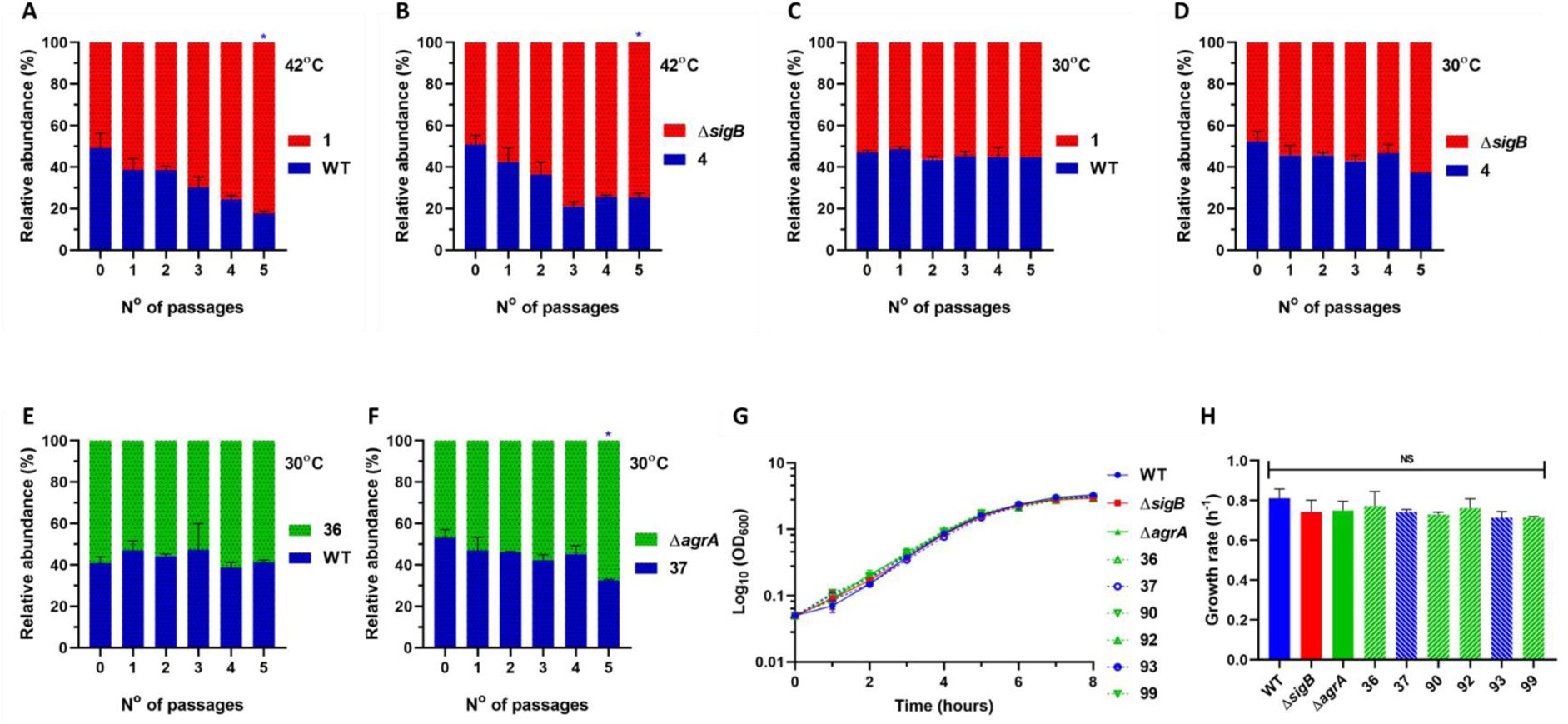
The competitive advantage conferred by *sigB* and *agr* alleles is limited at 30°C. Competitions experiments were performed with white versus grey colonies isolated during the *in vitro* evolution experiments. (A, B) Show competitions made at 42°C and (C, D, E, F) were made at 30°C. Blue bars represent the wild type and white coloured strains, while red and green represent *sigB*^−^ and *agr*^−^, respectively. (G) Growth curves were performed at 30°C in BHI using wild type, Δ*sigB*, Δ*agrA* and colonies isolated during the *in vitro* evolution experiment at 30°C. (H) Growth rates were calculated from growth curves from (G). All competitions were initiated with a 1:1 ratio of white:grey colonies. Data were generated from two independent biological replicates. Statistical analysis was performed using a paired student *t*-test in comparison to the passage 0 (*, *p*-value of <0.05).

**Figure S3 –.**
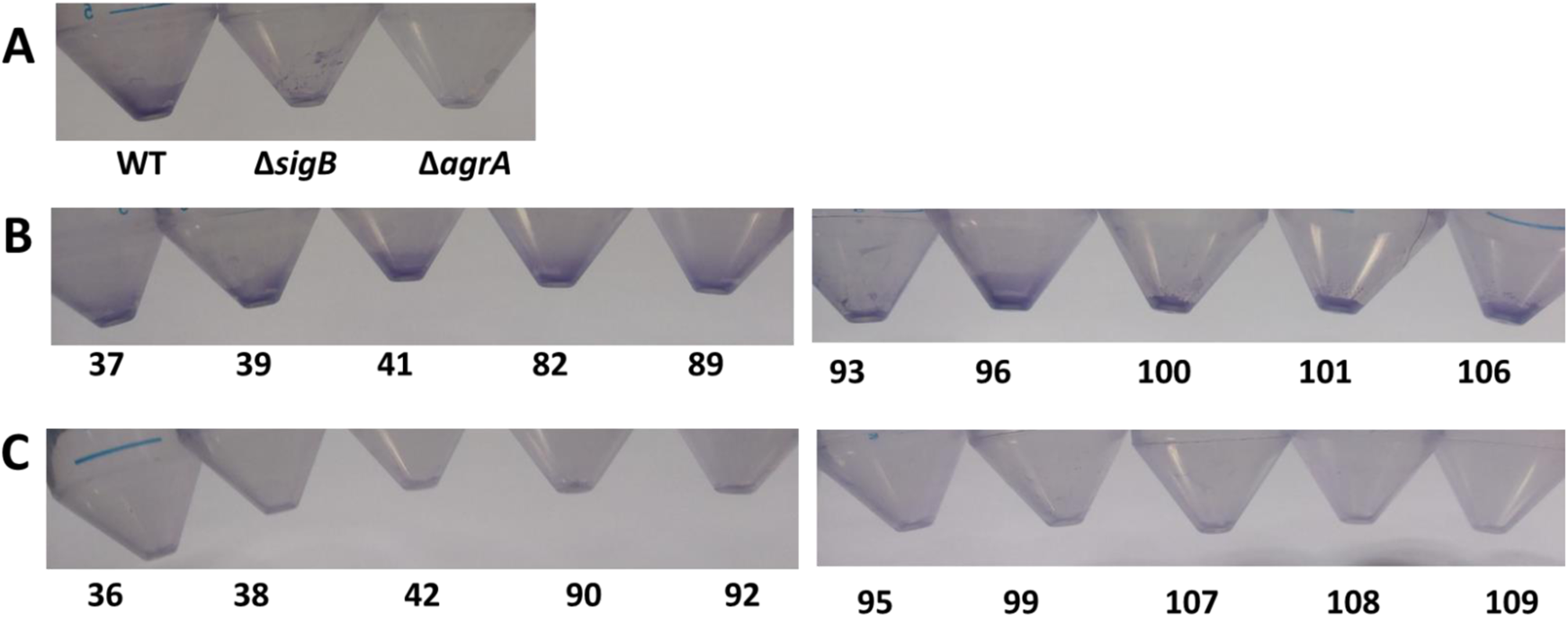
Grey colonies carrying *agr* alleles do not form biofilm in shaking conical tubes. Biofilm formation of wild type, Δ*sigB* and Δ*agrA* (A), phenotypically white (B) and grey (C) coloured isolates grown for 24 hours at 30°C with a constant agitation of 150 rpm. Pictures are representative of at least two independent biological replicates.

## References

1. Arnaud, M., Chastanet, A., & Débarbouillé, M. (2004). New Vector for Efficient Allelic Replacement in Naturally Nontransformable, Low-GC-Content, Gram-Positive Bacteria. Applied and Environmental Microbiology, 70(11), 6887–6891. https://doi.org/10.1128/AEM.70.11.6887-6891.2004

2. Asakura, H., Kawamoto, K., Okada, Y., Kasuga, F., Makino, S., Yamamoto, S., & Igimi, S. (2012). Intrahost passage alters SigB-dependent acid resistance and host cell-associated kinetics of Listeria monocytogenes. Infection, Genetics and Evolution, 12(1), 94–101. https://doi.org/10.1016/j.meegid.2011.10.014

3. Becker, L. A., Çetin, M. S., Hutkins, R. W., & Benson, A. K. (1998). Identification of the Gene Encoding the Alternative Sigma Factor ςB from Listeria monocytogenesand Its Role in Osmotolerance. Journal of Bacteriology, 180(17), 4547–4554. https://doi.org/10.1128/JB.180.17.4547-4554.1998

4. Boura, M., Keating, C., Royet, K., Paudyal, R., O’Donoghue, B., O’Byrne, C. P., & Karatzas, K. A. G. (2016). Loss of SigB in Listeria monocytogenes Strains EGD-e and 10403S Confers Hyperresistance to Hydrogen Peroxide in Stationary Phase under Aerobic Conditions. Applied and Environmental Microbiology, 82(15), 4584–4591. https://doi.org/10.1128/AEM.00709-16

5. Brøndsted, L., Kallipolitis, B. H., Ingmer, H., & Knöchel, S. (2003). KdpE and a putative RsbQ homologue contribute to growth of Listeria monocytogenes at high osmolarity and low temperature. FEMS Microbiology Letters, 219(2), 233–239. https://doi.org/10.1016/S0378-1097(03)00052-1

6. Cao, M., Bitar, A. P., & Marquis, H. (2007). A mariner-Based Transposition System for Listeria monocytogenes. Applied and Environmental Microbiology, 73(8), 2758–2761. https://doi.org/10.1128/AEM.02844-06

7. Chaturongakul, S., & Boor, K. J. (2004). RsbT and RsbV Contribute to σB-Dependent Survival under Environmental, Energy, and Intracellular Stress Conditions in Listeria monocytogenes. Applied and Environmental Microbiology, 70(9), 5349–5356. https://doi.org/10.1128/AEM.70.9.5349-5356.2004

8. Connolly, J. P. R., Roe, A. J., & O’Boyle, N. (2021). Prokaryotic life finds a way: Insights from evolutionary experimentation in bacteria. Critical Reviews in Microbiology, 47(1), 126–140. https://doi.org/10.1080/1040841X.2020.1854172

9. Deatherage, D. E., & Barrick, J. E. (2014). Identification of Mutations in Laboratory-Evolved Microbes from Next-Generation Sequencing Data Using breseq. In L. Sun & W. Shou (Eds.), Engineering and Analyzing Multicellular Systems: Methods and Protocols (pp. 165–188). Springer. https://doi.org/10.1007/978-1-4939-0554-6_12

10. den Dunnen, J. T., Dalgleish, R., Maglott, D. R., Hart, R. K., Greenblatt, M. S., McGowan-Jordan, J., Roux, A.-F., Smith, T., Antonarakis, S. E., & Taschner, P. E. M. (2016). HGVS Recommendations for the Description of Sequence Variants: 2016 Update. Human Mutation, 37(6), 564–569. https://doi.org/10.1002/humu.22981

11. Dorey, A., Marinho, C., Piveteau, P., & O’Byrne, C. (2019). Chapter One—Role and regulation of the stress activated sigma factor sigma B (σB) in the saprophytic and host-associated life stages of Listeria monocytogenes. In G. M. Gadd & S. Sariaslani (Eds.), Advances in Applied Microbiology (Vol. 106, pp. 1–48). Academic Press. https://doi.org/10.1016/bs.aambs.2018.11.001

12. Ferreira, A., Gray, M., Wiedmann, M., & Boor, K. J. (2004). Comparative Genomic Analysis of the sigB Operon in Listeria monocytogenes and in Other Gram-Positive Bacteria. Current Microbiology, 48(1), 39–46. https://doi.org/10.1007/s00284-003-4020-x

13. Fowler, V. G., Jr., Sakoulas, G., McIntyre, L. M., Meka, V. G., Arbeit, R. D., Cabell, C. H., Stryjewski, M. E., Eliopoulos, G. M., Barth Reller, L., Ralph Corey, G., Jones, T., Lucindo, N., Yeaman, M. R., & Bayer, A. S. (2004). Persistent Bacteremia Due to Methicillin-Resistant Staphylococcus aureus Infection Is Associated with agr Dysfunction and Low-Level In Vitro Resistance to Thrombin-Induced Platelet Microbicidal Protein. The Journal of Infectious Diseases, 190(6), 1140–1149. https://doi.org/10.1086/423145

14. Gaballa, A., Guariglia-Oropeza, V., Wiedmann, M., & Boor, K. J. (2019). Cross Talk between SigB and PrfA in Listeria monocytogenes Facilitates Transitions between Extra- and Intracellular Environments. Microbiology and Molecular Biology Reviews: MMBR, 83(4). https://doi.org/10.1128/MMBR.00034-19

15. García-Contreras, R., Nuñez-López, L., Jasso-Chávez, R., Kwan, B. W., Belmont, J. A., Rangel-Vega, A., Maeda, T., & Wood, T. K. (2015). Quorum sensing enhancement of the stress response promotes resistance to quorum quenching and prevents social cheating. The ISME Journal, 9(1), 115–125. https://doi.org/10.1038/ismej.2014.98

16. Garmyn, D., Augagneur, Y., Gal, L., Vivant, A.-L., & Piveteau, P. (2012). Listeria monocytogenes Differential Transcriptome Analysis Reveals Temperature-Dependent Agr Regulation and Suggests Overlaps with Other Regulons. PLOS ONE, 7(9), e43154. https://doi.org/10.1371/journal.pone.0043154

17. George, S. E., Hrubesch, J., Breuing, I., Vetter, N., Korn, N., Hennemann, K., Bleul, L., Willmann, M., Ebner, P., Götz, F., & Wolz, C. (2019). Oxidative stress drives the selection of quorum sensing mutants in the Staphylococcus aureus population. Proceedings of the National Academy of Sciences, 116(38), 19145–19154. https://doi.org/10.1073/pnas.1902752116

18. Gray, J., Chandry, P. S., Kaur, M., Kocharunchitt, C., Fanning, S., Bowman, J. P., & Fox, E. M. (2021). Colonisation dynamics of Listeria monocytogenes strains isolated from food production environments. Scientific Reports, 11(1), 12195. https://doi.org/10.1038/s41598-021-91503-w

19. Guerreiro, D. N., Arcari, T., & O’Byrne, C. P. (2020). The σB-Mediated General Stress Response of Listeria monocytogenes: Life and Death Decision Making in a Pathogen. Frontiers in Microbiology, 11. https://doi.org/10.3389/fmicb.2020.01505

20. Guerreiro, D. N., Wu, J., Dessaux, C., Oliveira, A. H., Tiensuu, T., Gudynaite, D., Marinho, C. M., Boyd, A., García-Del Portillo, F., Johansson, J., & O’Byrne, C. P. (2020). Mild Stress Conditions during Laboratory Culture Promote the Proliferation of Mutations That Negatively Affect Sigma B Activity in Listeria monocytogenes. Journal of Bacteriology, 202(9). https://doi.org/10.1128/JB.00751-19

21. Guldimann, C., Guariglia-Oropeza, V., Harrand, S., Kent, D., Boor, K. J., & Wiedmann, M. (2017). Stochastic and Differential Activation of σB and PrfA in Listeria monocytogenes at the Single Cell Level under Different Environmental Stress Conditions. Frontiers in Microbiology, 8, 348. https://doi.org/10.3389/fmicb.2017.00348

22. Hafner, L., Pichon, M., Burucoa, C., Nusser, S. H. A., Moura, A., Garcia-Garcera, M., & Lecuit, M. (2021). Listeria monocytogenes faecal carriage is common and depends on the gut microbiota. Nature Communications, 12(1), 6826. https://doi.org/10.1038/s41467-021-27069-y

23. Hall, B. G., Acar, H., Nandipati, A., & Barlow, M. (2014). Growth Rates Made Easy. Molecular Biology and Evolution, 31(1), 232–238. https://doi.org/10.1093/molbev/mst187

24. He, L., Le, K. Y., Khan, B. A., Nguyen, T. H., Hunt, R. L., Bae, J. S., Kabat, J., Zheng, Y., Cheung, G. Y. C., Li, M., & Otto, M. (2019). Resistance to leukocytes ties benefits of quorum sensing dysfunctionality to biofilm infection. Nature Microbiology, 4(7), 1114–1119. https://doi.org/10.1038/s41564-019-0413-x

25. Ireton, K., & Cossart, P. (1997). Host-Pathogen Interactions During Entry and Actin-Based Movement of Listeria Monocytogenes. Annual Review of Genetics, 31(1), 113–138. https://doi.org/10.1146/annurev.genet.31.1.113

26. Kim, H., Marquis, H., & Boor, K. J. (2005). ΣB contributes to Listeria monocytogenes invasion by controlling expression of inlA and inlB. Microbiology (Reading, England), 151(Pt 10), 3215–3222. https://doi.org/10.1099/mic.0.28070-0

27. Kumar, S., Parvathi, A., George, J., Krohne, G., Karunasagar, I., & Karunasagar, I. (2009). A study on the effects of some laboratory-derived genetic mutations on biofilm formation by Listeria monocytogenes. World Journal of Microbiology and Biotechnology, 25(3), 527–531. https://doi.org/10.1007/s11274-008-9919-8

28. Larsen, M. H., Kallipolitis, B. H., Christiansen, J. K., Olsen, J. E., & Ingmer, H. (2006). The response regulator ResD modulates virulence gene expression in response to carbohydrates in Listeria monocytogenes. Molecular Microbiology, 61(6), 1622–1635. https://doi.org/10.1111/j.1365-2958.2006.05328.x

29. Lauderdale, K. J., Boles, B. R., Cheung, A. L., & Horswill, A. R. (2009). Interconnections between Sigma B, agr, and Proteolytic Activity in Staphylococcus aureus Biofilm Maturation. Infection and Immunity, 77(4), 1623–1635. https://doi.org/10.1128/IAI.01036-08

30. Lecuit, M., Vandormael-Pournin, S., Lefort, J., Huerre, M., Gounon, P., Dupuy, C., Babinet, C., & Cossart, P. (2001). A Transgenic Model for Listeriosis: Role of Internalin in Crossing the Intestinal Barrier. Science. https://www.science.org/doi/abs/10.1126/science.1059852

31. Lee, Y.-J., & Wang, C. (2020). Links between S-adenosylmethionine and Agr-based quorum sensing for biofilm development in Listeria monocytogenes EGD-e. MicrobiologyOpen, 9(5), e1015. https://doi.org/10.1002/mbo3.1015

32. Liu, Y., Orsi, R. H., Gaballa, A., Wiedmann, M., Boor, K. J., & Guariglia-Oropeza, V. (2019). Systematic review of the Listeria monocytogenes σB regulon supports a role in stress response, virulence and metabolism. Future Microbiology, 14, 801–828. https://doi.org/10.2217/fmb-2019-0072

33. Marinho, C. M., Dos Santos, P. T., Kallipolitis, B. H., Johansson, J., Ignatov, D., Guerreiro, D. N., Piveteau, P., & O’Byrne, C. P. (2019). The σB-dependent regulatory sRNA Rli47 represses isoleucine biosynthesis in Listeria monocytogenes through a direct interaction with the ilvA transcript. RNA Biology, 16(10), 1424–1437. https://doi.org/10.1080/15476286.2019.1632776

34. Marinho, C. M., Garmyn, D., Gal, L., Brunhede, M. Z., O’Byrne, C., & Piveteau, P. (2020). Investigation of the roles of AgrA and σB regulators in Listeria monocytogenes adaptation to roots and soil. FEMS Microbiology Letters, 367(3). https://doi.org/10.1093/femsle/fnaa036

35. Mengaud, J., Ohayon, H., Gounon, P., Mège, R.-M., & Cossart, P. (1996). E-Cadherin Is the Receptor for Internalin, a Surface Protein Required for Entry of L. monocytogenes into Epithelial Cells. Cell, 84(6), 923–932. https://doi.org/10.1016/S0092-8674(00)81070-3

36. Monk, I. R., Gahan, C. G. M., & Hill, C. (2008). Tools for Functional Postgenomic Analysis of Listeria monocytogenes. Applied and Environmental Microbiology, 74(13), 3921–3934. https://doi.org/10.1128/AEM.00314-08

37. NicAogáin, K., & O’Byrne, C. P. (2016). The Role of Stress and Stress Adaptations in Determining the Fate of the Bacterial Pathogen Listeria monocytogenes in the Food Chain. Frontiers in Microbiology, 7, 1865. https://doi.org/10.3389/fmicb.2016.01865

38. O’Byrne, C. P., & Karatzas, K. A. G. (2008). Advances in Applied Microbiology (Vol. 65). Academic Press.

39. O’Donoghue, B. (2016). A Molecular Genetic Investigation into Stress Sensing in the Food-Borne Pathogen Listeria monocytogenes: Roles for RsbR and its Paralogues [Doctoral Thesis]. National University of Ireland, Galway.

40. O’Donoghue, B., NicAogáin, K., Bennett, C., Conneely, A., Tiensuu, T., Johansson, J., & O’Byrne, C. (2016). Blue-Light Inhibition of Listeria monocytogenes Growth Is Mediated by Reactive Oxygen Species and Is Influenced by σB and the Blue-Light Sensor Lmo0799. Applied and Environmental Microbiology, 82(13), 4017–4027. https://doi.org/10.1128/AEM.00685-16

41. Oliveira, A. H., Tiensuu, T., Guerreiro, D. N., Tükenmez, H., Dessaux, C., García-del Portillo, F., O’Byrne, C., & Johansson, J. (2021). Listeria monocytogenes requires the RsbX protein to prevent SigB-activation under non-stressed conditions. *Journal of Bacteriology*, JB.00486–21. https://doi.org/10.1128/JB.00486-21

42. Periasamy, S., Joo, H.-S., Duong, A. C., Bach, T.-H. L., Tan, V. Y., Chatterjee, S. S., Cheung, G. Y. C., & Otto, M. (2012). How Staphylococcus aureus biofilms develop their characteristic structure. Proceedings of the National Academy of Sciences, 109(4), 1281–1286. https://doi.org/10.1073/pnas.1115006109

43. Pfaffl, M. W. (2001). A new mathematical model for relative quantification in real-time RT–PCR. Nucleic Acids Research, 29(9), e45–e45. https://doi.org/10.1093/nar/29.9.e45

44. Pfaffl, M. W., Georgieva, T. M., Georgiev, I. P., Ontsouka, E., Hageleit, M., & Blum, J. W. (2002). Real-time RT-PCR quantification of insulin-like growth factor (IGF)-1, IGF-1 receptor, IGF-2, IGF-2 receptor, insulin receptor, growth hormone receptor, IGF-binding proteins 1, 2 and 3 in the bovine species. Domestic Animal Endocrinology, 22(2), 91–102. https://doi.org/10.1016/S0739-7240(01)00128-X

45. Quereda, J. J., Pucciarelli, M. G., Botello-Morte, L., Calvo, E., Carvalho, F., Bouchier, C., Vieira, A., Mariscotti, J. F., Chakraborty, T., Cossart, P., Hain, T., Cabanes, D., & García-del Portillo, F. 2013. (2013). Occurrence of mutations impairing sigma factor B (SigB) function upon inactivation of Listeria monocytogenes genes encoding surface proteins. Microbiology, 159(Pt_7), 1328–1339. https://doi.org/10.1099/mic.0.067744-0

46. Riedel, C. U., Monk, I. R., Casey, P. G., Waidmann, M. S., Gahan, C. G. M., & Hill, C. (2009). AgrD-dependent quorum sensing affects biofilm formation, invasion, virulence and global gene expression profiles in Listeria monocytogenes. Molecular Microbiology, 71(5), 1177–1189. https://doi.org/10.1111/j.1365-2958.2008.06589.x

47. Rieu, A., Lemaître, J.-P., Guzzo, J., & Piveteau, P. (2008). Interactions in dual species biofilms between Listeria monocytogenes EGD-e and several strains of Staphylococcus aureus. International Journal of Food Microbiology, 126(1), 76–82. https://doi.org/10.1016/j.ijfoodmicro.2008.05.006

48. Rieu, A., Weidmann, S., Garmyn, D., Piveteau, P., & Guzzo, J. (2007). agr System of Listeria monocytogenes EGD-e: Role in Adherence and Differential Expression Pattern. Applied and Environmental Microbiology, 73(19), 6125–6133. https://doi.org/10.1128/AEM.00608-07

49. Robinson, T., Smith, P., Alberts, E. R., Colussi-Pelaez, M., & Schuster, M. (2020). Cooperation and Cheating through a Secreted Aminopeptidase in the Pseudomonas aeruginosa RpoS Response. MBio, 11(2), e03090–19. https://doi.org/10.1128/mBio.03090-19

50. Shen, Y., Naujokas, M., Park, M., & Ireton, K. (2000). InlB-Dependent Internalization of Listeria Is Mediated by the Met Receptor Tyrosine Kinase. Cell, 103(3), 501–510. https://doi.org/10.1016/S0092-8674(00)00141-0

51. Sleator, R. D., Watson, D., Hill, C., & Gahan, C. G. M. Y. 2009. (2009). The interaction between Listeria monocytogenes and the host gastrointestinal tract. Microbiology, 155(8), 2463–2475. https://doi.org/10.1099/mic.0.030205-0

52. Spira, B., de Almeida Toledo, R., Maharjan, R. P., & Ferenci, T. (2011). The uncertain consequences of transferring bacterial strains between laboratories— RpoSinstability as an example. BMC Microbiology, 11(1), 248. https://doi.org/10.1186/1471-2180-11-248

53. Tiensuu, T., Andersson, C., Rydén, P., & Johansson, J. (2013). Cycles of light and dark co-ordinate reversible colony differentiation in Listeria monocytogenes. Molecular Microbiology, 87(4), 909–924. https://doi.org/10.1111/mmi.12140

54. Tiensuu, T., Guerreiro, D. N., Oliveira, A. H., O’Byrne, C., & Johansson, J. (2019). Flick of a switch: Regulatory mechanisms allowing Listeria monocytogenes to transition from a saprophyte to a killer. Microbiology, 165(8), 819–833. https://doi.org/10.1099/mic.0.000808

55. Traber, K. E., Lee, E., Benson, S., Corrigan, R., Cantera, M., Shopsin, B., & Novick, R. P. (2008). Agr function in clinical Staphylococcus aureus isolates. Microbiology (Reading, England), 154(Pt 8), 2265–2274. https://doi.org/10.1099/mic.0.2007/011874-0

56. Utratna, M., Cosgrave, E., Baustian, C., Ceredig, R., & O’Byrne, C. (2012). Development and optimization of an EGFP-based reporter for measuring the general stress response in Listeria monocytogenes. Bioengineered, 3(2), 93–103. https://doi.org/10.4161/bbug.19476

57. Vuong, C., Kocianova, S., Yao, Y., Carmody, A. B., & Otto, M. (2004). Increased Colonization of Indwelling Medical Devices by Quorum-Sensing Mutants of Staphylococcus epidermidis In Vivo. The Journal of Infectious Diseases, 190(8), 1498–1505. https://doi.org/10.1086/424487

58. Whiteley, M., Diggle, S. P., & Greenberg, E. P. (2017). Progress in and promise of bacterial quorum sensing research. Nature, 551(7680), 313–320. https://doi.org/10.1038/nature24624

59. WHO. (2018). *Listeriosis*. https://www.who.int/news-room/fact-sheets/detail/listeriosis

60. Williams, T., Bauer, S., Beier, D., & Kuhn, M. (2005). Construction and Characterization of Listeria monocytogenes Mutants with In-Frame Deletions in the Response Regulator Genes Identified in the Genome Sequence. Infection and Immunity. https://doi.org/10.1128/IAI.73.5.3152-3159.2005

61. Xia, Y., Xin, Y., Li, X., & Fang, W. (2016). To Modulate Survival under Secondary Stress Conditions, Listeria monocytogenes 10403S Employs RsbX To Downregulate σB Activity in the Poststress Recovery Stage or Stationary Phase. Applied and Environmental Microbiology, 82(4), 1126–1135. https://doi.org/10.1128/AEM.03218-15

62. Yuan, G., & Wong, S. L. (1995). Regulation of groE expression in Bacillus subtilis: The involvement of the sigma A-like promoter and the roles of the inverted repeat sequence (CIRCE). Journal of Bacteriology, 177(19), 5427–5433. https://doi.org/10.1128/jb.177.19.5427-5433.1995

63. Zetzmann, M., Sánchez-Kopper, A., Waidmann, M. S., Blombach, B., & Riedel, C. U. (2016). Identification of the agr Peptide of Listeria monocytogenes. Frontiers in Microbiology, 7, 989. https://doi.org/10.3389/fmicb.2016.00989

